# Fighting cancer with oncolytic viral therapy: identifying threshold parameters for success

**DOI:** 10.1101/2021.07.19.452846

**Authors:** Sana Jahedi, Lin Wang, James Watmough

## Abstract

We model interactions between cancer cells and free virus during oncolytic viral therapy. One of our main goals is to identify parameter regions that yield treatment failure or success. We show that the tumor size under therapy at a particular time is less than the size without therapy. Our analysis shows there are two thresholds for the horizontal transmission rate: a “Control threshold”, the threshold above which treatment is efficient, and an “optimum threshold”, the threshold beyond which infection prevalence reaches 100% and the tumor shrinks to its smallest size. Moreover, we explain how changes in the virulence level of the free virus alter the optimum threshold and the minimum tumor size. We identify a threshold for the virulence level of the virus and show how this threshold depends on the timescale of virus dynamics. Our results suggest that when the timescale of virus dynamics is fast, the administration of a more virulent virus leads to more tumor reduction. Conversely, when the viral timescale is slow, a higher virulence will have drawbacks on the results, such as high amplitude oscillations. Furthermore, our numerical observation depicts fast and slow dynamics. Our numerical simulations indicate there exists a two-dimensional globally attracting surface that includes the unstable manifold of the interior equilibrium. All solutions with positive initial conditions rapidly approach this two-dimensional attracting surface. In contrast, the trajectories on the attracting surface slowly tend to the periodic solution.

**Highlights:**

- The assumption that the viral load is in a quasi-steady state is relaxed, and infected cells are assumed to be mitotic.
- Our model strongly suggests that the tumor size is always reduced by therapy.
- We identify minimum tumor size, control threshold, and optimum threshold of the therapy.
- Our analysis shows optimal virulence level of oncolytic virus depends on the time scale of virus dynamics.
- When virus dynamic is slow, highly virulent virus causes long remission before relapse.

## 1. Introduction

The idea of using viruses as a cancer treatment was formed when doctors noticed that some cancer patients experienced a short period of remission after a viral infection [1]. Oncolytic viruses are natural or modified viruses that specifically target cancerous cells, replicate in them and eventually kill the infected cancer cells via lysis. Oncolytic viruses can also be armed with some cytokines or chemokines to activate antitumor immune cells [2, 3, 4, 5, 6], but here we only focus on their use as oncolytic agents.

To date, clinical trials conducted by using various oncolytic viruses have shown that these agents are less toxic than conventional therapies [7, 8], which makes this therapy popular. However, the therapeutic success of an oncolytic virus is limited [9, 10], and the complete eradication of a tumor has rarely been achieved when an oncolytic virus has been used as a single agent [11, 12]. This motivates us to study interactions between virus and cancer cells during oncolytic viral therapy to identify barriers and find ways to overcome them. Here, we use mathematical modeling to address the following questions: what are the outcomes of oncolytic viral therapy? Under which condition is a specific outcome possible? Can the oncolytic virus be used as a cure or only as a containment method? And how does the virulence of the virus affect the outcome of the therapy? Moreover, we examine how to optimize the efficiency of viral therapy by identifying which parameter regions will lead to the best outcome under the therapy.

Several mathematical models of oncolytic virus dynamics have been developed [13, 14, 15, 16, 17]. Komarova and Wodarz [18] studied the interactions between infected and uninfected cancer cells under the assumption that the turnover of the free virus is fast enough to maintain a quasi-steady state. They showed there is an optimum value for the death rate of infected cells at which the tumor reaches a minimum size. Long-term oscillatory behavior has not been observed in their model, but in vivo studies of oncolytic viruses have suggested oscillatory behavior [9]. This encourages us to consider a three-dimensional model by relaxing the quasi-steady state assumption, to capture all possible dynamics during an oncolytic viral therapy.

Similar to many viruses, oncolytic viruses come in two types: enveloped and naked. Some example of naked oncolytic viruses are adenovirus [19] and reovirus [20]. For examples of enveloped oncolytic viruses we can refer to vesicular stomatitis virus [21], type 1 herpes simplex virus [22], and Zika virus [23]. Enveloped oncolytic viruses can be easier for the host immune cells to remove than naked oncolytic viruses [24]. New virions of enveloped viruses can leave the membrane of their host without lysis, but naked viruses only leave the host cells via lysis. Hence, enveloped oncolytic viruses seem to be less efficient than naked oncolytic viruses. Our model describes the interaction of a naked oncolytic virus with infected and uninfected cancer cells during oncolytic viral therapy.

A virus can affect cell death mechanisms in various ways. A type of death that an infection can induce on a cell is lysis. When a cell becomes infected, the virus makes copies of itself until the infected cell’s membrane is burst, and new virions are released into the extracellular area. Most viruses are equipped with anti-apoptosis proteins that allow them to complete the viral reproduction cycle before an infected cell dies [25]. However, sometimes an infected cell dies quickly in response to an infection to prevent infection spread; the virions released in this way have not completed the viral reproduction cycle and are not infectious. This paper’s model describes the dynamics of oncolytic viral therapy when viruses can complete the viral reproduction cycle before an infected cell dies.

Similar to Komarova and Wodarz [18], we consider using a mitotic virus. In other words, there are two pathways for transmission of infection: horizontal transmission, where an uninfected cell encounters free virus and becomes infected and vertical transmission, where infected cells produce infected daughter cells through mitosis.

In some cancers, larger tumors have an increased risk of metastasis. Koscielny et al. [26] showed that the incidence of metastasis increases as the size of a breast tumor increases. In a study on the survival experience of 1894 patients with invasive breast tumors, Narod [27] showed that an increase in tumor size decreases the 15-year survival rate of patients. Hence, the size to which the tumor shrinks is a crucial factor in identifying how successful cancer treatment is. A therapy that causes considerable tumor shrinkage may be used as a tumor control method and, in combination with other therapies, may lead to tumor eradication. This inspires us to investigate how a virus should be engineered to shrink the steady-state of the tumor size compared to the case that no treatment occurs. It is essential to know for which area in parameter space tumor size is minimum. As we will show, there exists a threshold for the horizontal rate of infectivity above which the tumor size reaches its minimum.

To optimize the therapy, one needs to know how virulent the virus should be. Virulence is the negative impact caused by a viral infection on the functionality of host cells [28], such as a lower cell division rate or a higher mortality rate after infection. Here, we consider using an aggressive virus and explicitly include a rate associated with the effect of infection on cancer cell viability into the model. Our goal is to assess how the rate related to virulence affects final tumor size. Recent studies suggest that extreme levels of viral aggression are not recommended [18]. Here, we show that when the virus becomes more aggressive, the system shows oscillatory behavior. We derive a threshold for virulence above which a periodic solution exists. Our numerical observations suggest this periodic solution is globally stable: that is to say, it attracts every solution that starts from a positive point which is not on the stable manifold of the positive equilibrium.

Capturing fast and slow dynamics is very beneficial to understand the treatment outcome in the longer term, especially when the fast behavior is followed immediately by a slow behavior. For example, when dynamics are fast, and the treatment outcome is followed only for a short time, it may sound that the treatment is successful in the sense that huge tumor shrinkage is observed on the fast time scale, but tumors may appear again on the slower time scale. On the other hand, when the dynamics are too slow, tumor growth may not be noticeable when solutions are only observed for a short time, but in practice, a tumor may get big enough to be a concern again within some years. We show for which parameter region and for which state spaces fast and slow dynamics occur.

This paper has a new mathematical approach to Hopf-bifurcation and attractors that can be of clinical importance. We describe attracting periodic orbits that arise in Hopf bifurcations of our model. There is a steady-state solution *E*(*m*) where *m* is a real-valued bifurcation parameter. When a stable steady-state bifurcates to a stable periodic solution as *m* is changed, the traditional Hopf approach focuses on the new periodic solution. Our approach is to focus on the sometimes fast-slow nature of Hopf bifurcations. Our model is three-dimensional. Near the Hopf bifurcation point, the fixed point’s two-dimensional unstable manifold is bounded by the stable periodic orbit that is created at the Hopf bifurcation. There is also a transverse contracting direction. Linearizing at the fixed point and computing the eigenvalues, there is a negative eigenvalue *A*(*m*) *<* 0 and a pair of complex eigenvalues whose real part is *r*(*m*). Following the creation of the periodic orbits, *r*(*m*) is small and positive. An initial condition near the fixed point will approach the unstable manifold very quickly and then will oscillate with growing amplitude approaching the limit cycle very slowly. Figure 1 shows the behavior of this paper’s main treatment model. It is essential for the clinician to focus on the entire period of growth instead of just the limit cycle, when the tumor takes years to approach a limit cycle.

**Figure 1:**
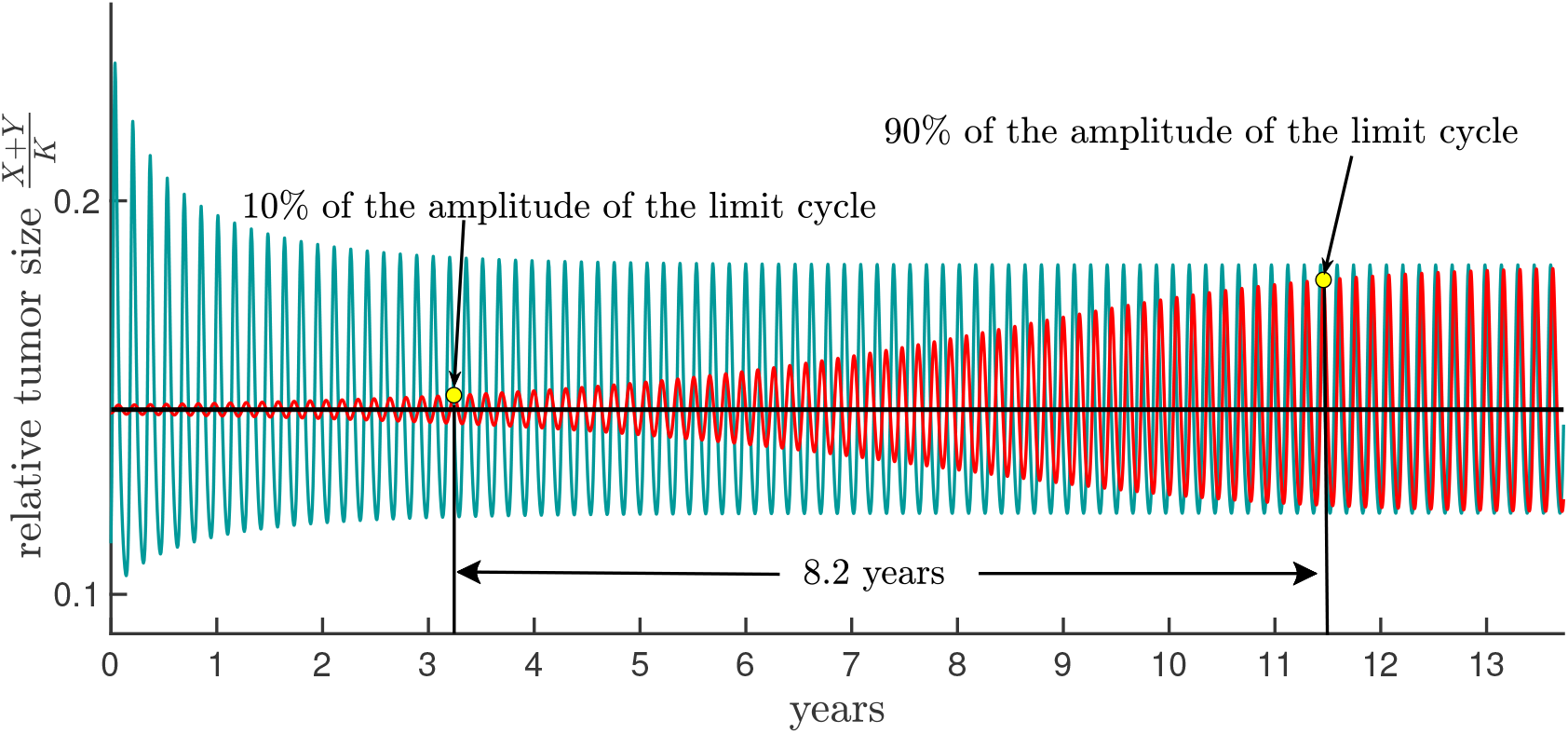
Slow dynamics. The black curve is the equilibrium of Model (1) that loses its stability and bifurcates in a stable limit cycle when the Hopf bifurcation occurs. As the figure suggests, the red and green curves tend very slowly to the attracting limit cycle. The time between peaks is approximately 57 days. The red curve takes 8.2 years to grow from 10% of the amplitude of the limit cycle to 90%, a very long time for a cancer patient; see the text for the parameter values.

There is a second aspect that makes this slow growth an important clinical factor. Hopf bifurcations often occur in pairs. Figure 2 shows a pair of Hopf bifurcations of our model. In Figure 1 *m* is 0.1, where *r*(*m*) is near its maximum value between the bifurcation values *m*_1_ and *m*_2_. Note that an oscillatory trajectory’s amplitude has an exponential growth rate that slows as the trajectory moves away from equilibrium for the generic or typical Hopf bifurcation. Hence, for a fixed value of *m* such as *m* = 0.1, *r*(*m*) is an upper bound on the exponential growth rate of the amplitude of trajectories on the unstable manifold for a generic Hopf-bifurcation. In Figure 1, *r*(*m* = 0.1) = 0.00099 ∼ 0.001, which means the amplitude grows by a factor of about one part in a thousand per day. This rate is only slightly smaller than the maximum value 0.0011 of *r*(*m*), which occurs at *m* = 0.086. From the clinical point of view, it is best to view such unstable manifolds as consisting of a disk of nearly periodic orbits. The patient may not live long enough to see the limit cycle.

**Figure 2:**
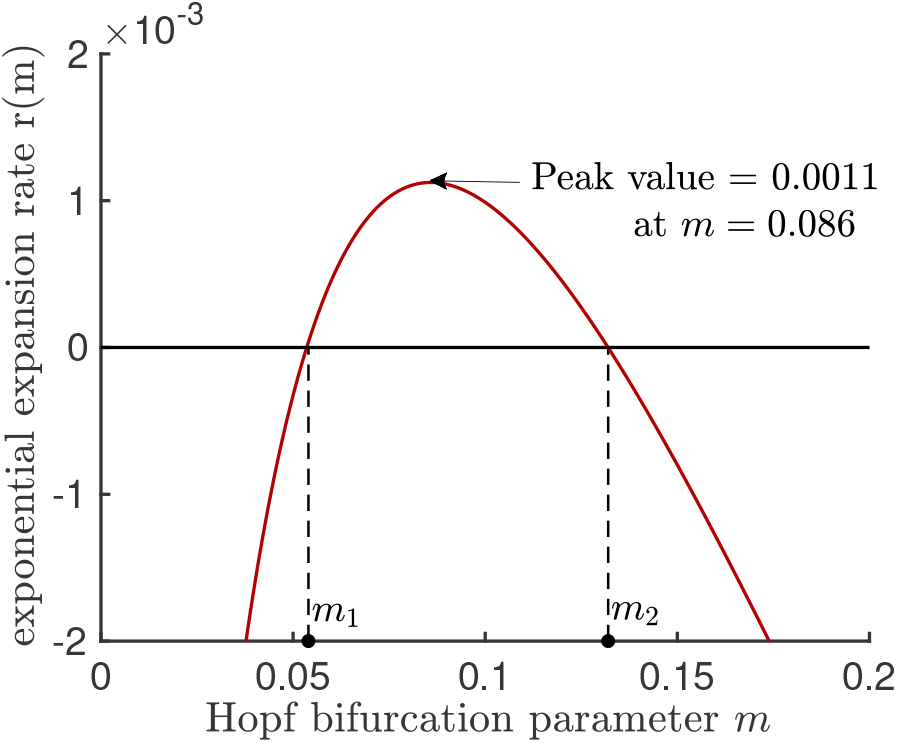
The exponential growth rate *r*(*m*). In our model there are two Hopf bifurcations from the equilibrium *E*_*p*_(*m*) as the parameter *m* varies, namely *m*_1_ and *m*_2_. Linearizing the dynamics at the fixed point *E*_*p*_(*m*), there is a pair of complex eigenvalues whose real part is *r*(*m*). Note that *r*(*m*_1_) = *r*(*m*_2_) = 0.

## 2. Model and Assumptions

We make the following assumptions to achieve a tractable model that helps us focus on our research questions mentioned earlier.

A1: To model nutrient limitations, we assume the mitosis rates are density dependent, dropping linearly to zero at a maximal tumor size *K*. We denote the per capita mitosis rates of uninfected and infected cancer cells at zero density by 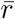 and 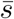, respectively.

A2: As we mentioned before, we assume there are two pathways for transmission of infection: *β* is the horizontal rate of infectivity per virus and 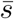, the mitosis rate of infected cancer cells, is the vertical transmission rate, see Figure 3.

**Figure 3:**
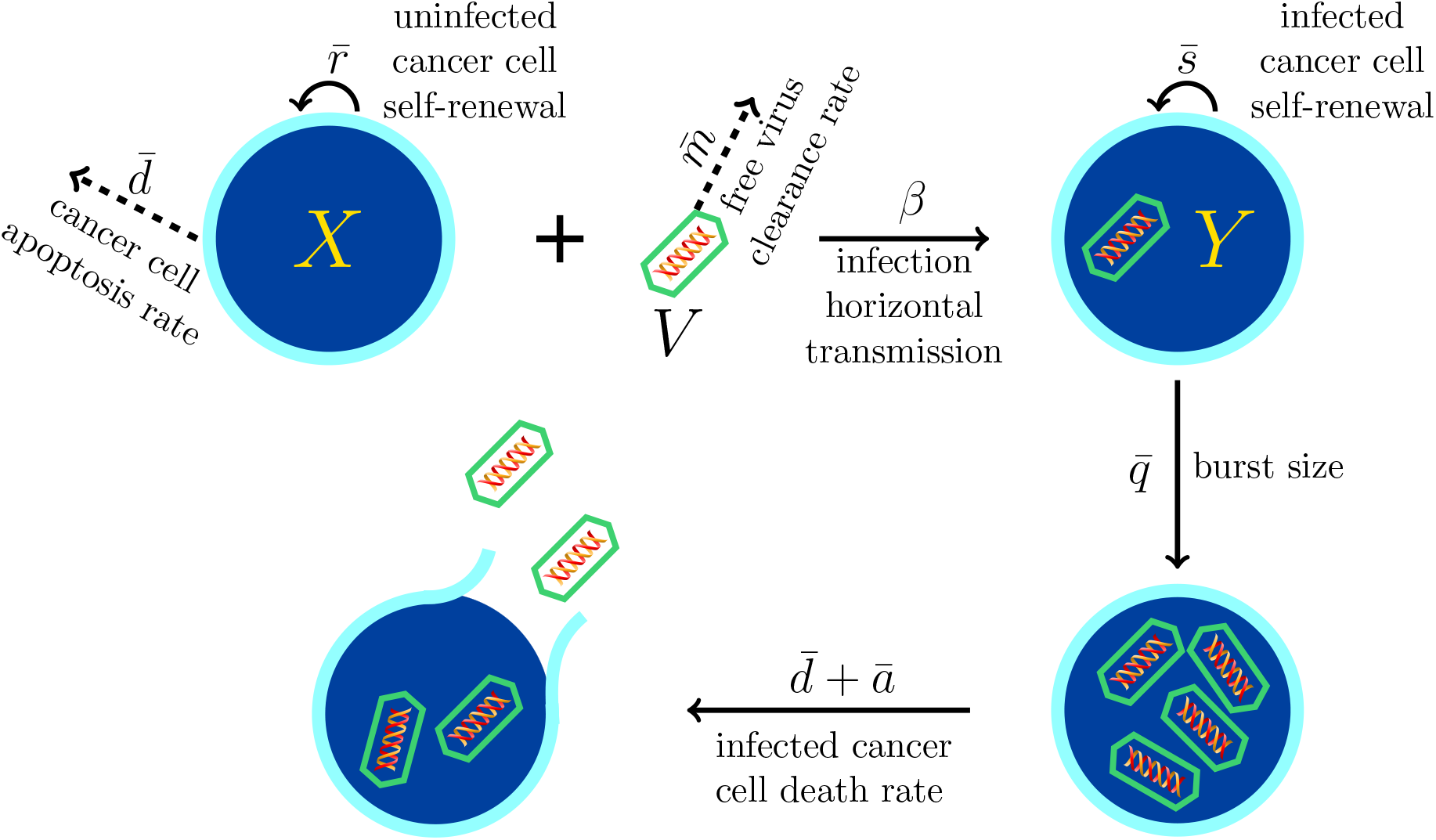
A schematic description of the Model (1). *X, Y*, and *V* denote population densities of uninfected cancer cells, infected cancer cells, and the free virus, respectively.

A3: Furthermore, we assume the vertical transmission is perfect. Meaning an infected cell never produces an uninfected daughter cell.

A4: Uninfected and infected cancer cells die at rates 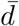 and 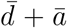 respectively. These rates are assumed to be lower than the per capita mitosis rates 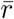 and 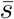, respectively.

A5: The virus is virulent, meaning

- infection reduces cell growth; the mitosis rate of infected cancer cells, 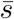, is lower than mitosis rate of uninfected cancer cells, 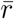, i.e. 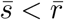.
- infection elevates apoptosis; parameter ā > 0, denotes the elevation in death rate of cancer cells after infection.

A6: As we mentioned in the introduction, we assume our model shows the dynamics of a virus that can complete the viral reproduction cycle before the host infected cell dies. We further assume that no virion can leave the membrane of an infected cell before the infected cell dies. As Figure 3 illustrates, the free virus makes 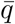 copies of itself in each infected cell, and the infected cell dies at the rate 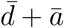. Therefore, 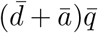 new virions are produced per each dead infected cell.

A7: Here in the interest of keeping consistent with the earlier models [18, 29, 30] that we built our model on, we neglect removal of the free viruses with infection.

Under the above assumptions we have the following model:

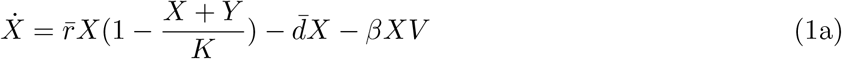

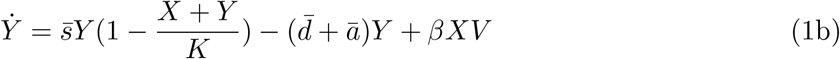

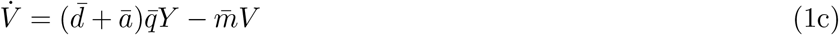

where *X, Y*, and *V* denote the population density of uninfected cancer cells, infected cancer cells, and free virus, respectively. 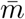 in Eq. (1c) is the clearance rate of the free virus, and finally, *X* + *Y* denotes the tumor size. Figure 3 illustrates the viral cycle. First, viral incidence occurs between an uninfected cancer cell *X* and the free virus *V* at rate *β*. Once the free virus penetrates the target cell, the target cells become infected. Then the free virus replicates itself till the membrane of the infected cell is disrupted and the new virions are released in the extracellular space.

As we mentioned in assumption (A6), Model (1) demonstrates the dynamics of a naked oncolytic virus. Therefore, it is assumed that new virions only are released to the extracellular area after the infected cell dies. Therefore, the rate at which new virions are reproduced is 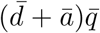. However, when using an enveloped virus, new virions can leave the membrane of an infected cell without lysis. For Model (1) to describe dynamics of an oncolytic virus, one should change equation 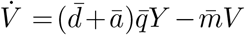 to 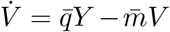 (in this case 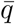 denotes the rate at which new virions are produced by each infected cell). Under such a change, mathematically, there will be no difference in how we analyze the results. Since we report the results for the rescaled model. For both scenarios, with different rescalings, the same dimensionless model can be obtained. However, if we extend Model (1) to consider interactions with virus-specific immune responses, depending on which types of virus-specific immune response we consider, the outcome of therapy may be different depending on which kind of oncolytic virus (naked vs. enveloped) is considered.

To simplify our analysis, we rescale the parameters and variables of Model (1) as follows:

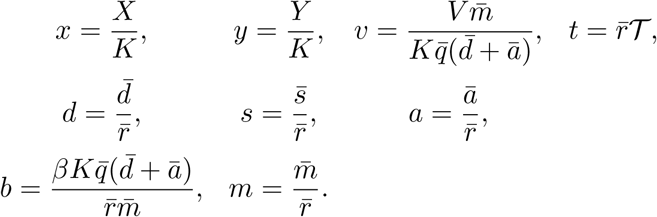

Also for simplification we introduce two parameter combinations *X*_*U*_ = 1 −*d* and *Y*_*I*_ = 1 −(*d* +*a*)*/s*. After applying these changes Model (1) turns to the following dimensionless model, the algebraic calculations are provided in Appendix A.

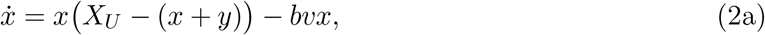

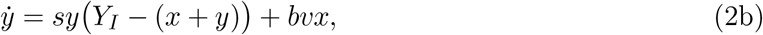

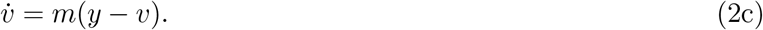

Notice that when *m* is a large (small) number according to Eq. 2c, 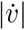 is large (small, respectively). Therefore, throughout this work, we use parameter *m* as a measure of the time scale of virus dynamics.

Assumptions (A4) and (A5) in the terms of new parameters can be restated as follows;

(H1): *s <* 1; this holds since from assumption (A5), 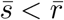, hence 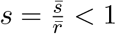.

(H2): *X*_*U*_ *> Y*_*I*_ *>* 0; these hold because according to assumption (A4), 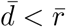 and 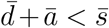. Hence, 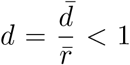 and 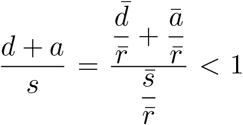. Thus, *X*_*U*_ *>* 0 and *Y*_*I*_ *>* 0. According to assumption (H1), *s <* 1. Therefore, *sd <* 1(*d* + *a*). Hence, 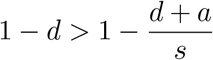, meaning *X*_*U*_ *> Y*_*I*_.

When modeling a biological phenomenon, since the variables represent population densities, we should show the solutions always stay non-negative and bounded. The following lemma says each solution of Model (2) starting in 𝒲 will remain in 𝒲. The shaded wedge shape area in Figure 4(a) represents 𝒲. Since 𝒲 is a bounded subset of the first octant, therefore solutions of Model (2) are bounded and non-negative.

**Figure 4:**
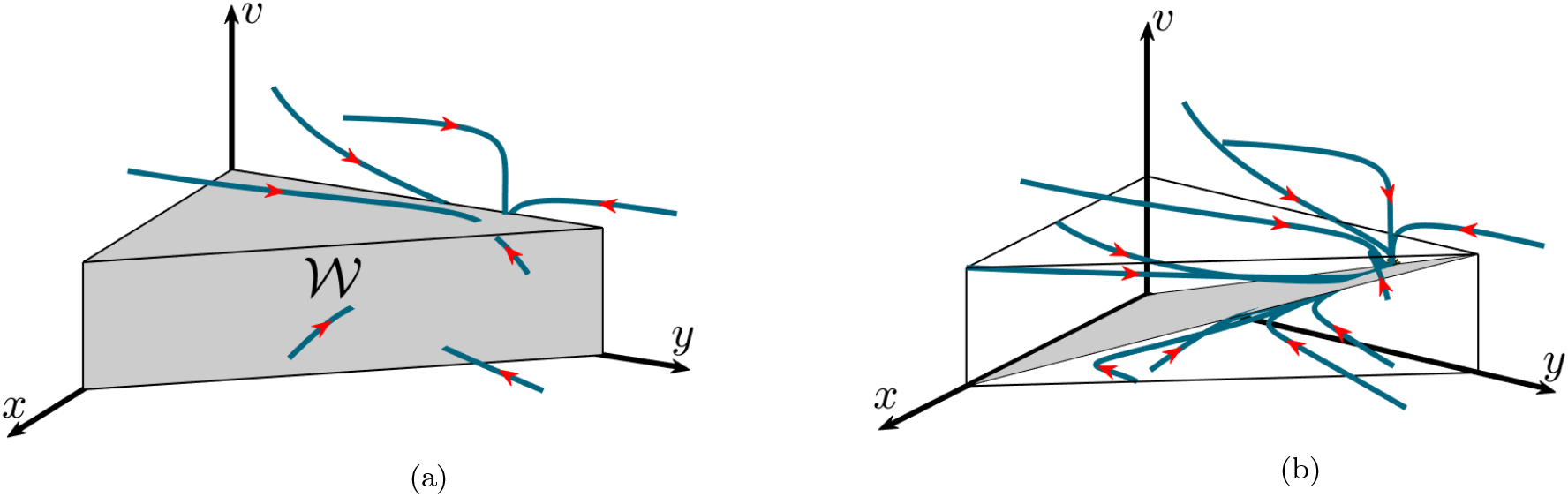
𝒲 is positively invariant and globally attractive. (a) This plot demonstrates that the trajectory of each solution that starts from a point in the first octant eventually gets attracted to 𝒲. (b) This plot is the same as plot (a) but includes the *y* = *v* plane’s intersection with 𝒲. When a solution starts above this plane, *y < v* so *v* decreases, and when a solution starts below this plane, *v* increases.

### Lemma 1.

For Model (2) the set

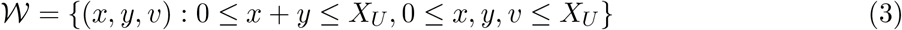

is positively invariant and globally attractive.

**Proof**. To show 𝒲 is positively invariant, it is enough to show a trajectory that starts from a point on the boundaries of 𝒲 either remains on the boundaries of 𝒲 for all *t* ≥ 0 or it tends inward 𝒲. Each solution of Model (2) that starts with an initial point on the boundaries of 𝒲 which is not located on the *v* axis, *x* axis or on the *y* − *v* plane, points inward of 𝒲. Each solution that starts from an initial point on the boundaries of 𝒲, which is located on the *v* axis, the *x* axis or *y* − *v* plane tends to (0, 0, 0), (*X*_*U*_, 0, 0) or (0, *Y*_*I*_, *Y*_*I*_), respectively (for details see Appendix B). (0, 0, 0), (*X*_*U*_, 0, 0) and (0, *Y*_*I*_, *Y*_*I*_) are all in boundaries of 𝒲. Therefore, 𝒲 is positively invariant.

When *x* + *y* ≥ *X*_*U*_ and *x, y >* 0, Eqs. (2a) and (2b) yield the following equation.

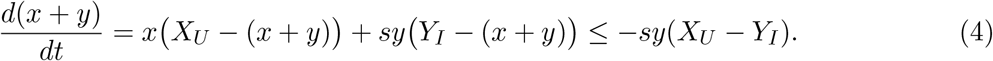

The above equation suggests if *x* + *y* ≥ *X*_*U*_ and *x, y >* 0, then 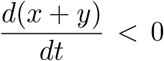. In other words, trajectory of each solution starting from a point (*x*(0), *y*(0), *v*(0)) that *x*(0) + *y*(0) ≥ *X*_*U*_ and *x*(0), *y*(0) *>* 0, moves in the direction of decreasing *x* + *y*, therefore, it tends inward 𝒲.

When *v* ≥ *X*_*U*_ and *v > y*, then 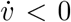. Hence *v* decreases and also any solution starting on the *x* +*y* = *X*_*U*_ plane points towards wedge in the direction of decreasing *x* +*y*. Therefore each solution starting from an initial point (*x*(0), *y*(0), *v*(0)), for which *x*(0) + *y*(0) *< X*_*U*_ and *v*(0) *> X*_*U*_, tends to a point in 𝒲.

Therefore, each solution with an initial point (*x*_0_, *y*_0_, *v*_0_) such that *x*_0_, *y*_0_, *v*_0_ ≥ 0 and (*x*_0_, *y*_0_, *v*_0_) ∉ 𝒲, eventually tends to a point in 𝒲.

Here after we only consider solutions with initial points in 𝒲. The shaded plane in the Figure 4(b) is the plane *y* = *v* intersection with 𝒲. All equilibria of Model (2) are on this plane.

The tumor size, meaning the total density of infected and uninfected cancer cells, determines whether the therapy is successful. The following proposition shows that the tumor size under oncolytic viral therapy always remains lower than the tumor size with no therapy over time.

Let 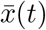, *t* ≥ 0, be the tumor size without therapy. That is, 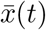 is the solution of Model (2) with initial condition *x*(0), *y*(0), *v*(0), where *x*(0) *>* 0, *y*(0) = *v*(0) = 0. Here *x*(*t*)+*y*(*t*), *t* ≥ 0 shows the tumor size with therapy. That is, the solution to Model (2) with initial condition (*x*(0), *y*(0), *v*(0)), where *x*(0) *>* 0, *v*(0) *>* 0, *y*(0) ≥ 0. Then we have the following.

### Proposition 1.

Under Model (2) assume the virus is introduced at time *t* = 0, either directly into cells or into the extracellular environment. Then the tumor size under therapy *x*(*t*) + *y*(*t*) at time *t >* 0 will be less than the tumor size without therapy 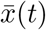. I.e. if 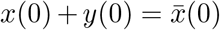 and *v*(0) *>* 0, *y*(0) ≥ 0, then 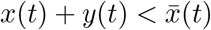 for all *t >* 0.

**Proof**. According to Eq. (2b), for each solution of Model (2) that starts with *x*(0) *>* 0 and *v*(0) *>* 0 and *y*(0) = 0, *y* variable is initially increasing 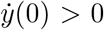. Hence, we conclude that *y*(*t*) *>* 0 for all *t >* 0. Now suppose the contrary assumption, that there exists *t*^*∗*^ *>* 0 such that 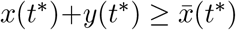, i.e.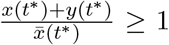. Define 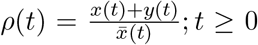. Notice that *ρ*(*t*) is a continuous function. Now we will show *ρ*(*t*) is strictly decreasing on the set of all *t* for which *ρ*(*t*) ≥ 1. Eqs. (2a) and (2b) yield to

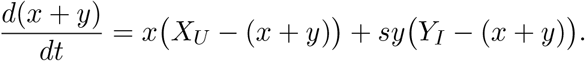

Since *y >* 0 for all *t >* 0, *Y*_*I*_ *< X*_*U*_, and *s <* 1,

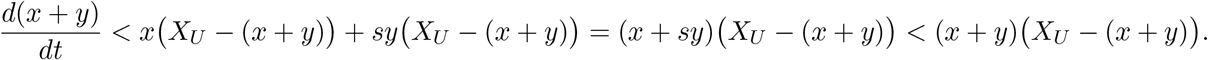

Since *x* + *y* ≠ 0,

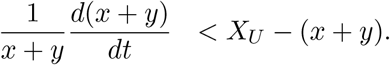

For all *t >* 0 such that *ρ*(*t*) ≥ 1 (i.e. 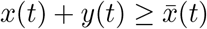),

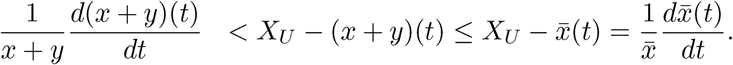

Hence, when 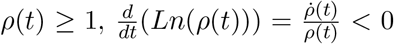. Therefore, *ρ*(*t*) is strictly decreasing on the set of all *t >* 0 for which *ρ*(*t*) ≥ 1. But *ρ*(*t*) is a continuous function and *ρ*(0) = 1. Therefore, *ρ* is strictly decreasing on the set of all *t* ≥ 0 for which *ρ*(*t*) ≥ 1. Therefore *ρ*(0) *> ρ*(*t*^*∗*^) ≥ 1, contradicting our assumption.

In the proof of the following proposition, we use comparison theorem [31], for convenience of the readers, the version of comparison theorem that we use is provided in Appendix C.1.

### Proposition 2.

(I) The tumor size under the therapy *x*(*t*) + *y*(*t*) will remain between *Y*_*I*_ and *X*_*U*_ if the initial tumor size is between *Y*_*I*_ and *X*_*U*_. I.e,

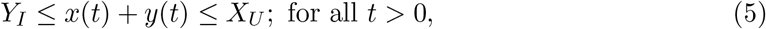

if *Y*_*I*_ ≤ *x*(0) + *y*(0) ≤ *X*_*U*_.

(II) For each solution (*x*(*t*), *y*(*t*), *v*(*t*)) with an initial condition (*x*(0), *y*(0), *v*(0)), where *x*(0) *>* 0, *y*(0) ≥ 0 there exists *t*^*∗*^ *>* 0 such that *Y*_*I*_ ≤ *x*(*t*^*∗*^) + *y*(*t*^*∗*^) ≤ *X*_*U*_.

**Proof of (I)**.

Eqs. (2a) and (2b) yield the following equation.

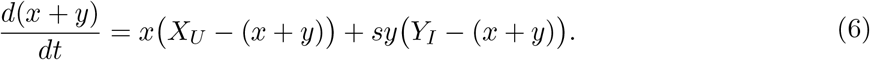

Since *x, y* ≥ 0, *Y*_*I*_ *< X*_*U*_ and *s <* 1 therefore, *sy(Y*_*I*_ − (*x* + *y*)) ≤ *y(X*_*U*_ − (*x* + *y*)). Hence,

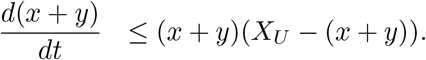

Therefore, if *x*(0) + *y*(0) ≤ *X*_*U*_, then according to the comparison theorem, *x*(*t*) + *y*(*t*) ≤ *X*_*U*_ for all *t* ≥ 0.

Similarly, using *s*(*x* + *y*) ≤ *x* + *sy* from Eq. (6)

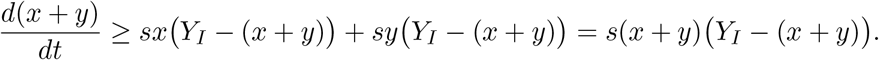

Therefore, if *x*(0) + *y*(0) ≥ *Y*_*I*_, then according to comparison theorem, *x*(*t*) + *y*(*t*) ≥ *Y*_*I*_ for all *t* ≥ 0.

**Proof of (II)**. Now assume (*x*(*t*), *y*(*t*), *v*(*t*)) is a solution with (*x*(0), *y*(0), *v*(0)), where *x*(0), *y*(0) *>* 0. We will show there exist *t*^*∗*^ *>* 0 such that *Y*_*I*_ ≤ *x*(*t*^*∗*^) + *y*(*t*^*∗*^) ≤ *X*_*U*_.

As we showed in the proof of Lemma 1, [0, *X*_*U*_] is globally attracting, so we only need to show that for each solution (*x*(*t*), *y*(*t*), *v*(*t*)) with initial condition *x*(0), *y*(0), *v*(0) where 0 *< x*(0) +*y*(0) *< Y*_*I*_ and *x*(0), *y*(0) *>* 0 there exists *t*^*∗*^ such that *x*(*t*^*∗*^) + *y*(*t*^*∗*^) ≥ *Y*_*I*_.

By contrary assume that the set *U* := {(*x*(*t*), *y*(*t*), *v*(*t*)) ∈ 𝒲: 0 *< x*(0) + *y*(0) *< Y*_*I*_, *x*(0), *y*(0) *>* 0} is positively invariant. If *x* + *y < Y*_*I*_ and *x, y >* 0 according to Eq. (6), 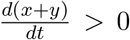, therefore, *x*(*t*) + *y*(*t*) is increasing along each solution (*x*(*t*), *y*(*t*), *v*(*t*)) ∈ *U* and it is bounded above by *Y*_*I*_. Hence, 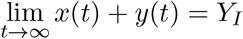, for any (*x*(*t*), *y*(*t*), *v*(*t*)) ∈ *U*. But *Y*_*I*_ is not in *U*, this contradicts by *U* being positively invariant.

## 3. Equilibria and their stability

Model (2) can have four equilibria in 𝒲, which we will refer to as follows: the tumor eradication equilibrium, *E*_0_ = (0, 0, 0), the treatment failure equilibrium, *E*_*U*_ = (*X*_*U*_, 0, 0), 100% infection prevalence equilibrium, *E*_*I*_ = (0, *Y*_*I*_, *Y*_*I*_) and the partially infected equilibrium, 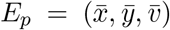, where

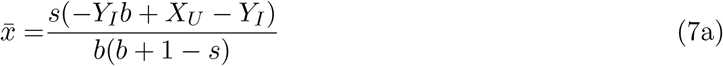

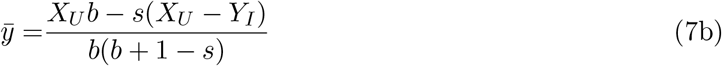

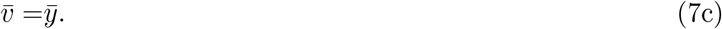

The following lemma tells under which conditions the above equilibria exist.

### Lemma 2.

*E*_0_, *E*_*U*_, *E*_*I*_ always exist. *E*_*p*_ is positive if and only if

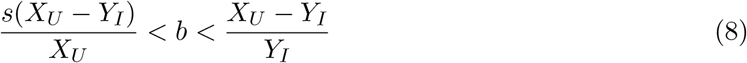

**Proof**. According to assumption (H2), *X*_*U*_ *>* 0 and *Y*_*I*_ *>* 0, thus *E*_*U*_ and *E*_*I*_ always exist. If 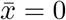, then simple calculations shows *E*_*p*_ = *E*_*I*_. Similarly, if 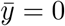, then *E*_*p*_ = *E*_*U*_. But, we are interested in the case that both 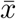 and 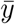 are positive. According to assumption (H1), *s <* 1. Thus, *b* + 1 − *s >* 0. Hence, *E*_*p*_ is positive if and only if both followings are satisfied:

1. − *Y*_*I*_*b* + *X*_*U*_ − *Y*_*I*_ *>* 0
2. *X*_*U*_*b* − s(*X*_*U*_ − *Y*_*I*_) *>* 0

Therefore, *E*_*p*_ is positive if and only if 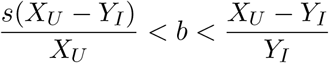.

We will refer to *E*_*p*_ as positive equilibrium. The following lemma summarizes the conditions under which each boundary equilibrium is locally asymptotically stable:

### Lemma 3.

1. *E*_0_ is always unstable
2. *E*_*U*_ = (*X*_*U*_, 0, 0) is locally stable if and only if 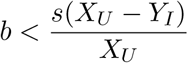.
3. *E*_*I*_ = (0, *Y, Y*) is locally stable if and only if 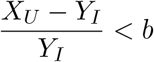

**Proof of (1)**. Eigenvalues of Jacobian of Model (2) at *E*_0_ are −*m, X*_*U*_, *sY*_*I*_. According to assumption (H2), *X*_*U*_, *Y*_*I*_ are both positive. Since *E*_0_ has two positive eigenvalues, so it is always unstable.

**Proof of (2)**. Jacobian of Model (2) at *E*_*U*_ always has a negative eigenvalue, −*X*_*U*_. The other two remaining eigenvalues are eigenvalues of the following matrix.

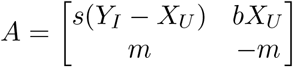

tr*A* = −*m* + *s*(*Y*_*I*_ − *X*_*U*_) and det 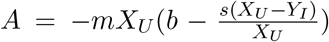. According to assumption (H2), *Y*_*I*_ *< X*_*U*_. Hence, *trA* is always negative. Therefore, the local stability of *E*_*U*_ is only determined by the sign of det *A*. Hence, *E*_*U*_ is locally stable if and only if det *A >* 0. Thus, *E*_*U*_ is locally stable if and only if 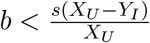.

**Proof of (3)**. Jacobian of Model (2) at equilibrium *E*_*I*_ has eigenvalues *λ*_*I*1_ = −*m, λ*_*I*2_ = −*sY*_*I*_, and *λ*_*I*3_ = *X*_*U*_ − *Y*_*I*_ − *bY*_*I*_. The first two eigenvalues are negative, so *E*_*I*_ is locally stable if and only if *λ*_*I*3_ is negative. In other words, *E*_*I*_ is locally stable if and only if 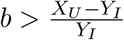

Proposition 2 and Lemma 3 together show when 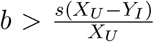 tumor shrinkage occurs and when 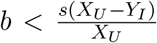 treatment fails. According to Lemma 3 when 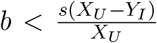, the treatment failure equilibrium *E*_*U*_ is stable and our numerical observations illustrate when an equilibrium is locally stable, it is also globally stable. I.e., each solution starting from a positive initial point will tend to that stable equilibrium. Therefore, when 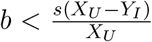, regardless of initial tumor size, administrating oncolytic virus has no effect, and the tumor grows to the maximum possible tumor size *X*_*U*_. Furthermore, Proposition 2 shows eventual tumor size is always smaller than or equal to *X*_*U*_, but Lemma 3 says when 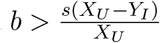, eventual tumor size is never *X*_*U*_. Hence, when 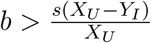 eventually tumor size is smaller than *X*_*U*_. Therefore, effective result by therapy will be observable when 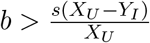. Hence, we call 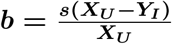 **tumor control threshold**.

Lemma 3 says when 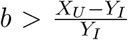, 100% infection prevalence equilibrium *E*_*I*_ is stable. In this case eventually tumor size is *Y*_*I*_. Additionally, Proposition 2 says *Y*_*I*_ is the optimum tumor size for a specific virus (fixed set of parameters), therefore we call 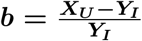 **optimum threshold** of *b* for therapy.

Figure 5 is a graphical representation of Proposition 2. This figure shows regardless of time scale of virus dynamics, tumor size under therapy remains between *Y*_*I*_ and *X*_*U*_. This figure shows tumor size under therapy decreases when the rescaled infectivity rate *b* increases. Notice that according to (H1), *s <* 1 and according to (H2), *Y*_*I*_ *< X*_*U*_, therefore 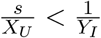, multiplying this inequality by (*X*_*U*_ − *Y*_*I*_) implies 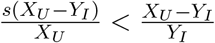. I.e, control threshold is always strictly lower than optimum threshold. Hence, Figure 5 is a generic bifurcation diagram of Model (2).

**Figure 5:**
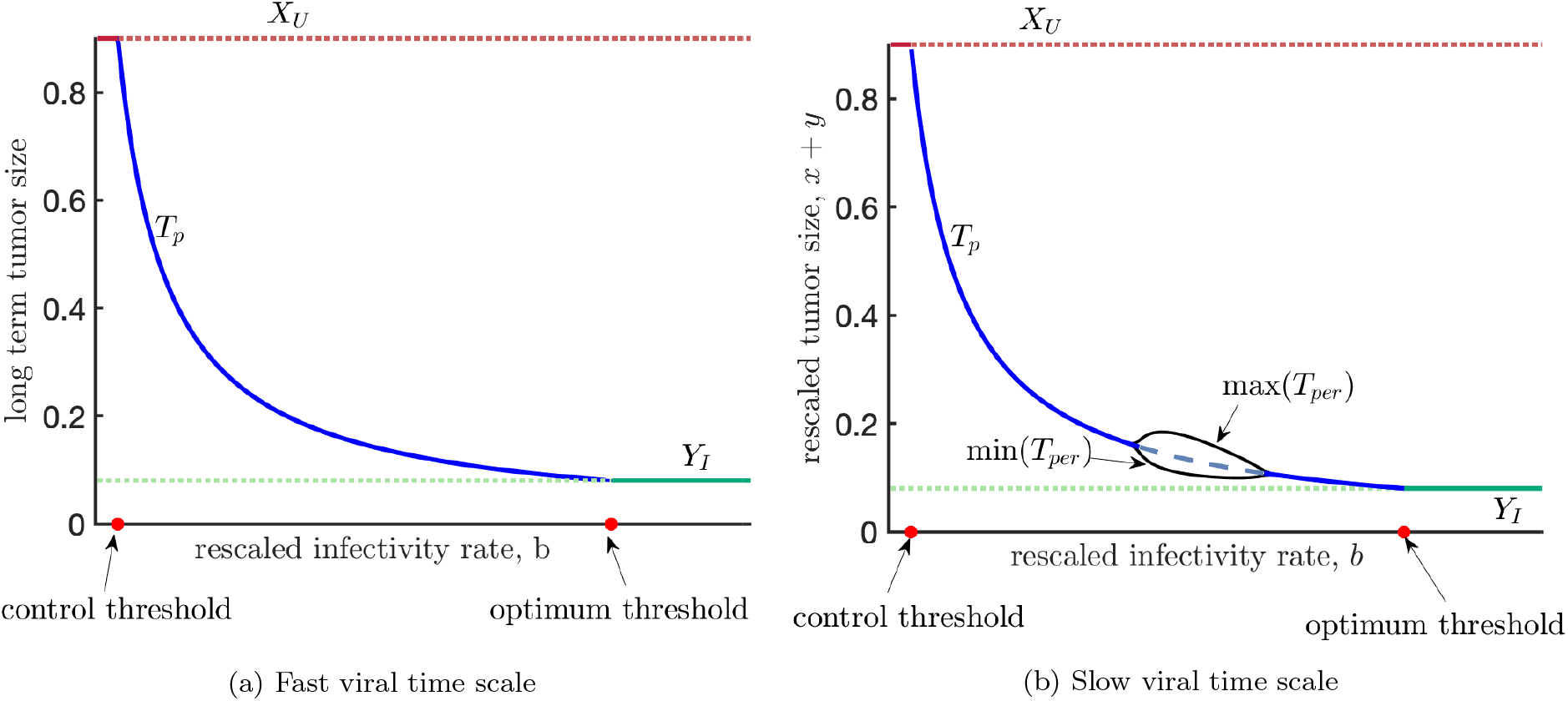
The outcome of therapy is enhanced when the horizontal rate of infection transmission increases. In both plots, maximum and minimum tumor size is sketched. When an equilibrium is globally stable, eventually, all the solutions will tend to it. Therefore, in that case, the maximum and minimum tumor size will be the same. As both plots represent, when *b* is smaller than the control threshold, treatment fails, and when *b* is greater than the optimum threshold, minimum tumor size by therapy *Y*_*I*_ is obtained. In both cases, the tumor size under therapy remains between *Y*_*I*_ and *X*_*U*_. When the viral time scale is fast, same as plot(a), no long-term oscillatory behavior will be observable, and when the viral time scale is slow, same as plot(b), sustained oscillation is observable. As plot (b) indicates, tumor size remains between *Y*_*I*_ and *X*_*U*_ even when sustained oscillations are observable. This simulation is conducted under the set of parameters *X*_*U*_ = 0.9, *Y*_*I*_ = 0.08, and *s* = 0.5. *m* = 1 in plot(a) and *m* = 0.1 in plot(b). In this plot, a dashed (solid) curve means that the corresponding equilibrium is unstable (stable) for that value of the parameter.

## 4. Hopf-bifurcation and sustained oscillations

A Hopf-bifurcation occurs in a system when by changes in one of the parameters of the system periodic solutions appear, i.e., in one side of the bifurcation point, there is no limit cycle, but as the bifurcation parameter passes through a bifurcation point, a periodic solution appears [32]. For the convenience of the reader, the Hopf bifurcation Theorem is provided in Appendix C. In this section, we will identify the parameter space at which Hopf bifurcation occurs.

Verifying the occurrence of a Hopf-bifurcation is often involved with the algebraic calculation of eigenvalues. In 1998, Liu [33] proposed a criterion for the occurrence of Hopf bifurcation without calculation of eigenvalues for a general system of ordinary differential equation of order *n*. Here, we use this criterion in the proof of Theorem 2. Hence, below we restate the criterion “Routh-Hopf” for a three-dimensional system of ordinary differential equations and provide a simpler proof for this special case.

Consider a three-dimensional system of ordinary differential equations is given, meaning

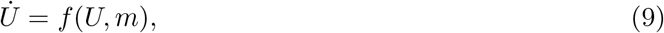

where *U* ∈ ℝ^3^ and *m* ∈ ℝ.

Assume (*U*_*eq*_(*m*), *m*) ∈ ℝ^3+1^ is a curve of stationary points for the system (9). Denote the characteristic polynomial of *Df* (*U*_*eq*_(*m*), *m*) by *P*_*m*_. Meaning

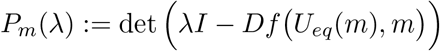

*P*_*m*_ is a degree three polynomial therefore, it can be written as *P*_*m*_(*λ*) = *λ*^3^ + *P*_2_(*m*)*λ*^2^ + *P*_1_(*m*)*λ* + *P*_0_(*m*).

Define Routh function as *H*(*m*) := *P*_2_(*m*)*P*_1_(*m*) − *P*_0_(*m*) [34].

### Theorem 1

(**Routh-Hopf Criterion)**. Assume that *P*_0_ ≠ 0. A Hopf bifurcation occurs at *m* = *m*^*∗*^ for the system (9) if the two following conditions are satisfied.

1. *H* (*m*\*) = 0,
2. 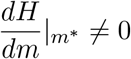

**Proof**. Let us assume that *P*_*m*_ has two complex roots *λ*_1_ = *a* + *bi, λ*_2_ = *a* − *bi* and a real root *c*, where *a* = *a*(*m*), *b* = *b*(*m*) and *c* = *c*(*m*). Hence, *P*_*m*_ can be written as

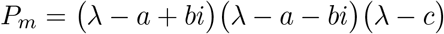

After simplification we have

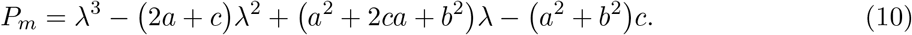

From Equation (10), Routh function n *H* = *P*_2_*P*_1_ −*P*_0_ can be written as *H* = −2*a* ·(*a*^2^ +2*ac* +*b*^2^ +*c*^2^). Therefore, *H* = −2*a* (*a* + *c*)^2^ + *b*^2^. Since *P*_0_ ≠ 0, therefore *H* is zero if and only if *a* = 0. Also, Since *P*_0_ ≠ 0, *a* and *b* cannot both be zero simultaneously.

Assume that *a*(*m*^*∗*^) = 0. We will show 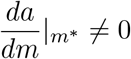 if and only if 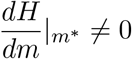.

As we showed above *H* = −2*a(*(*a* + *c*)^2^ + *b*^2)^, therefore

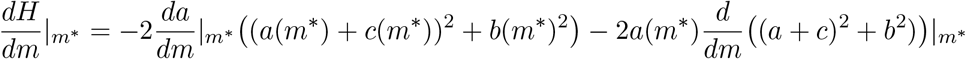

Since *a*(*m*^*∗*^) = 0, therefore

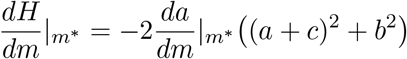

Since *P*_0_ ≠ 0, then (*a* + *c*)^2^ + *b*^2^ ≠ 0, therefore

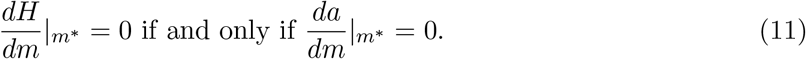

We have now shown (*i*) and (*ii*) imply the conditions of occurrence of a Hopf bifurcation (see Appendix C), so by Hopf bifurcation theorem, a Hopf-bifurcation occurs when (i) and (ii) hold.

### Theorem 2.

Let 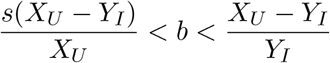. Define,

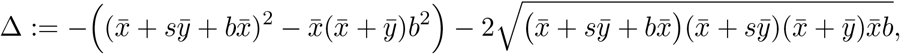

where 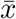 and 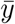 are same as Eq. (7). Then we have the following.

i. If Δ *<* 0, then *E*_*p*_ is a stable equilibrium.
ii. If Δ *>* 0, then two Hopf bifurcations occur as the parameter *m* changes. We denote these Hopf-bifurcation points by *m*_1_ and *m*_2_, where *m*_1_ *< m*_2_.
iii. If Δ *>* 0, then *E*_*p*_ is a stable equilibrium if and only if *m* ∈ (−∞, *m*_1_) ∪ (*m*_2_, ∞).

**Proof**. The Jacobian of Model (2) at 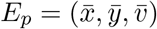 is

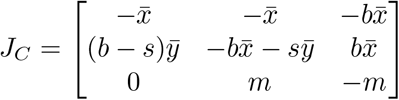

The characteristic equation for *J*_*C*_ is

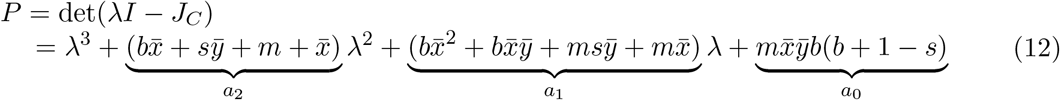

According to assumption (H1), *s <* 1. Therefore, *a*_1_, *a*_2_ and *a*_3_ are always positive. Routh function H for the above characteristic polynomial is as follows.

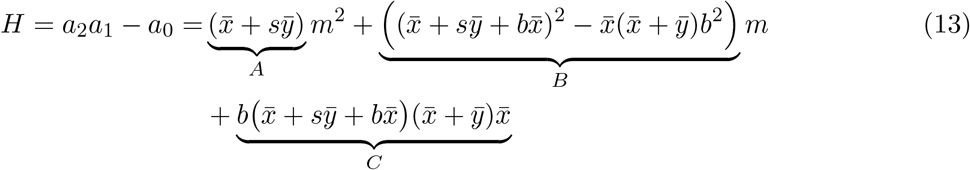

Since *A, C >* 0, *H* = 0 has positive solutions for *m* if and only if *B <* 0 and *B*^2^ −4*AC* ≥ 0. In other words, *H* has positive solutions for *m* if and only if 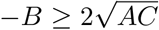. Notice that 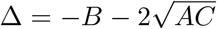. Hence, *H* = 0 only has solutions if and only if Δ ≥ 0.

When Δ *<* 0, *H* is always positive. As we mentioned above *a*_1_, *a*_2_ and *a*_3_ are also positive. Thus, according to Routh-Hurwitz stability criterion [34] equilibrium *E*_*p*_ is locally asymptotically stable. This proves part (i) of the theorem.

When Δ *>* 0, the quadratic *H* has two distinct positive roots *m*_1_ and *m*_2_, where *m*_1_ *< m*_2_. Of course, 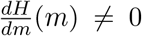 for *m* = *m*_1_, *m*_2_. Therefore according to Routh-Hopf Criterion, Hopf-bifurcations occur as *m* changes through *m*_1_ and *m*_2_. This proves part (ii) of the theorem.

When Δ *>* 0, and *m* ∈ (−∞, *m*_1_) ∪ (*m*_2_, ∞), then Routh function is positive, therefore according to Routh-Hurwitz criterion *E*_*p*_ is locally asymptotically stable. This proves part (iii).

Note that when Δ = 0, then the Routh function has a repeated root. Hence, there exists an *m*^*∗*^ such that *H*(*m*^*∗*^) = 0, but 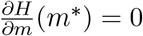. Thus, the conditions for the occurrence of Hopf bifurcation with respect to parameter *m* are not satisfied at *m* = *m*^*∗*^. Therefore, when Δ = 0, for any *m* ≠ *m*^*∗*^ *E*_*p*_ is an stable equilibrium and it is a non-hyperbolic equilibrium at *m* = *m*^*∗*^. In our numerical simulations, we observed that for *m* = *m*^*∗*^, the equilibrium *E*_*p*_ is globally stable, and every solution starting from a positive initial point tends to *E*_*p*_ very slowly, in the way that oscillating solutions are decaying very slowly, tending eventually to *E*_*p*_. Notice that when function *H* changes the sign, the real part of eigenvalues of *E*_*p*_ is changing the sign. Therefore, any bifurcation curve bifurcating from *E*_*p*_ should also cross *H* = 0. Since *H* is a function of all the parameters of Model (2), one may choose a different bifurcation parameter, here we have chosen *m* since we are interested in the effect of the time scale of virus dynamics on the outcome of therapy.

The following proposition allows us to choose the surface *y* − *v* = 0 as the Poincare section. This proposition tells that each positive point is either on the stable manifold of positive equilibrium *E*_*p*_ or its trajectory crosses *y* − *v* = 0 surface infinitely many times. There are situations where *E*_*p*_ appears to be a global attractor, attracting all the trajectories with positive initial values. Then, trajectories can spiral to *E*_*p*_, crossing the *y* = *v* surface infinitely many times.

### Proposition 3

(**Global dynamics)**. When 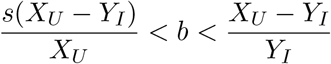, each trajectory starting from a positive point either crosses the surface *y* − *v* = 0 infinitely many times or its positive limit set is *E*_*p*_.

In the following proof we use notation *ω*_*p*_ to denote the forward limit set of a solution *p*(*t*) = (*x*^*p*^(*t*), *y*^*p*^(*t*), *v*^*p*^(*t*)) of Model (2). Strictly speaking, ω_*p*_ = {X ∈ 𝒲: ∃*t*_*n*_ ≥ 0 such that 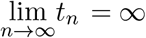 and 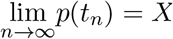 [35].

**Proof of Proposition 3**. Assume the trajectory *p*(*t*) = (*x*^*p*^(*t*), *y*^*p*^(*t*), *v*^*p*^(*t*)) does not cross the *y* = *v* plane in a finite time. Then it remains on one side of the plane *y* = *v* for all *t* ≥ 0. For specificity, let us assume *y*^*p*^(*t*) ≥ *v*^*p*^(*t*) for all *t* ≥ 0.

Call *U* := {(*x, y, v*) ∈ 𝒲; *y* ≥ *v*}. *U* is a compact set. Define *L*: ℝ^3^ → *R* such that *L*(*x, y, v*) = *v, L* is a continuous function and 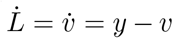, therefore *L*(*p*(*t*)) is monotonic (non-decreasing) for all *t* ≥ 0 and bounded subset of *U*. Therefore, lim_*t→∞*_ *L*(*p*(*t*)) exists. Assume *L*_0_ = lim_*t→∞*_ *L*(*p*(*t*)). Hence, lim_*t→∞*_ *v*^*p*^(*t*) = *L*_0_.

Suppose *X*(0) is in *ω*_*p*_. Since the forward limit set *ω*_*p*_ is positively invariant, *X*(*t*) ∈ *ω*_*p*_ for all *t* ≥ 0. For a fixed *t*^*∗*^ *>* 0, there exists a sequence *t*_*n*_ ≥ 0 such that 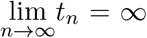 and 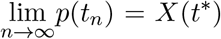. Since *L* is a continuous function 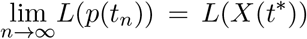. Hence, *L* (*X*(*t*\*)) = *L*_0_. Therefore, *L*(*X*(*t*)) = *L*_0_ for all *t* ≥ 0. Hence, 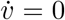 on *X*(*t*). Hence, *y* = *v* on *X* so 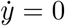. Since *v* = *y* = *L*_0_ are fixed on *X*(*t*), by solving Eq. (2b) for *x*(*t*), then there are two possibilities; either *L*_0_ = *Y*_*I*_, in this case *x* = 0, or *L*_0_ ≠ *Y*_*I*_, in this case 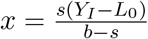. Hence, 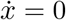 on *X*. So *X*(*t*) is an equilibrium. Since 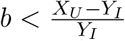, *E*_*I*_ cannot attract solutions with a positive initial condition, therefore, *L*_0_ ≠ *Y*_*I*_. Thus, *ω*_*p*_ = {*E*_*P*_} and *p*(*t*) goes to *ω*_*p*_ which equals *E*_*p*_.

We have proved that if a trajectory starting from a positive point does not cross surface *y* − *v* in a finite time, it will eventually tend to *E*_*p*_. If that trajectory crosses the surface *y* − *v* = 0 in a finite time, there are two options: either it crosses the surface *y* − *v* = 0 in a finite time or eventually tends to *E*_*p*_. Following the same pattern, we conclude that if a trajectory starting from a positive point does not tend to *E*_*p*_, it crosses the surface *y* − *v* = 0 infinitely many times.

## 5. Numerical observations

In this section, we report some observations that are illustrated numerically.

As it is mentioned above in Theorem 2 when Δ *>* 0 and *m*_1_ *< m < m*_2_ the equilibrium *E*_*p*_ is unstable and a locally stable limit cycle exists. In Figure 6 we have conducted a simulation such that a limit cycle exists: the black closed curve is the limit cycle, the black circle is equilibrium *E*_*p*_, and the brown curve passing through equilibrium *E*_*p*_ is its stable manifold. Our numerical observations suggest that when a limit cycle exists, it is almost everywhere globally stable. I.e., each solution starting from a positive initial point that is not located on the stable manifold of the equilibrium *E*_*p*_ tends to the limit cycle. In Figure 6, each solution starting from a point on the brown curve tends to *E*_*p*_. Each solution starting from a positive initial point which is not on the brown curve, tends to the limit cycle. Therefore, only some solutions (green and red curves) are sketched to have a clearer figure here. The red curves start from a solution outside of the limit cycle near the stable manifold of *E*_*p*_. They jump into the area enclosed by the limit cycle and spiral out from the nearby equilibrium *E*_*p*_, and tend to the limit cycle. The green curve oscillations’ amplitude gets smaller and spirals in towards the limit cycle. As the figure illustrates, the disk which is traced by the red curve, starting very nearby Ep and bounds to the limit cycle, is the unstable manifold of the equilibrium *E*_*p*_. The red disk, together with the equilibrium *E*_*p*_ and its boundary, the limit cycle forms a global attracting surface which we denote by G. Each solution with a positive initial condition tends to G.

**Figure 6:**
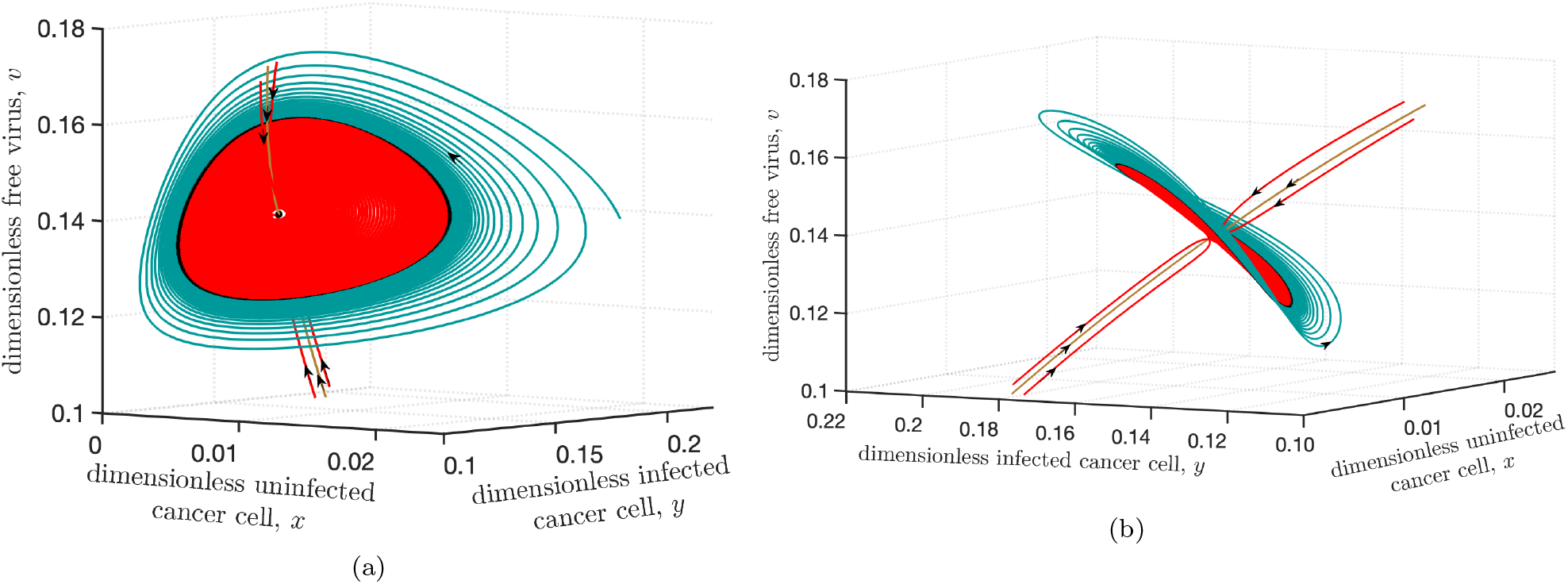
The global attracting surface. **(a)** The black curve represents a stable limit cycle, and the black circle is *E*_*p*_. Each solution starting from a positive point (except for the steady state *E*_*p*_ and its one dimensional stable manifold) approaches to this limit cycle. The red disk (traced out by the red curves) is the unstable manifold of *E*_*p*_, and the limit cycle is its boundary. Plot **(b)** is the same figure as plot (a), which has been rotated to show the stable manifold of equilibrium *E*_*p*_. The brown curve shows one dimensional stable manifold of *E*_*p*_. Every solution starting from a point on the brown curve gets attracted to the *E*_*p*_. The disk centered at equilibrium *E*_*p*_ with its boundary, the limit cycle, is (numerically) a global attracting surface. This simulation is conducted for 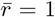, *s* = 0.5, *X*_*U*_ = 0.9, *Y*_*I*_ = 0.08, *m* = 0.1, and *b* ∼ 5.35. This *b* is the mid point of the interval 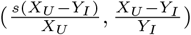, on which *E*_*p*_ exists.

Figure 7 shows bifurcation of tumor size with respect to changes in parameter *m* when 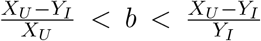. Our simulations show that when *m < m*_1_ or *m > m*_2_ the fixed point 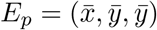 is globally stable, strictly speaking it attracts all the solutions starting from a positive initial point. Therefore, when *m < m*_1_ or *m > m*_2_, the steady state tumor size is 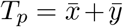. When *m*_1_ *< m < m*_2_, all the solutions starting from a positive point which is not on the stable manifold of *E*_*p*_ tends to the limit cycle, so in this case the tumor size is oscillating, so it has a maximum, denoted by the upper curve *max*(*T*_*per*_) and a minimum, denoted by the lower curve *min*(*T*_*per*_). As this figure illustrates the Hopf-bifurcation is global in the sense that as *m* changes a continuous path of periodic solutions sources from *m*_1_ and sinks in *m*_2_ [36].

**Figure 7:**
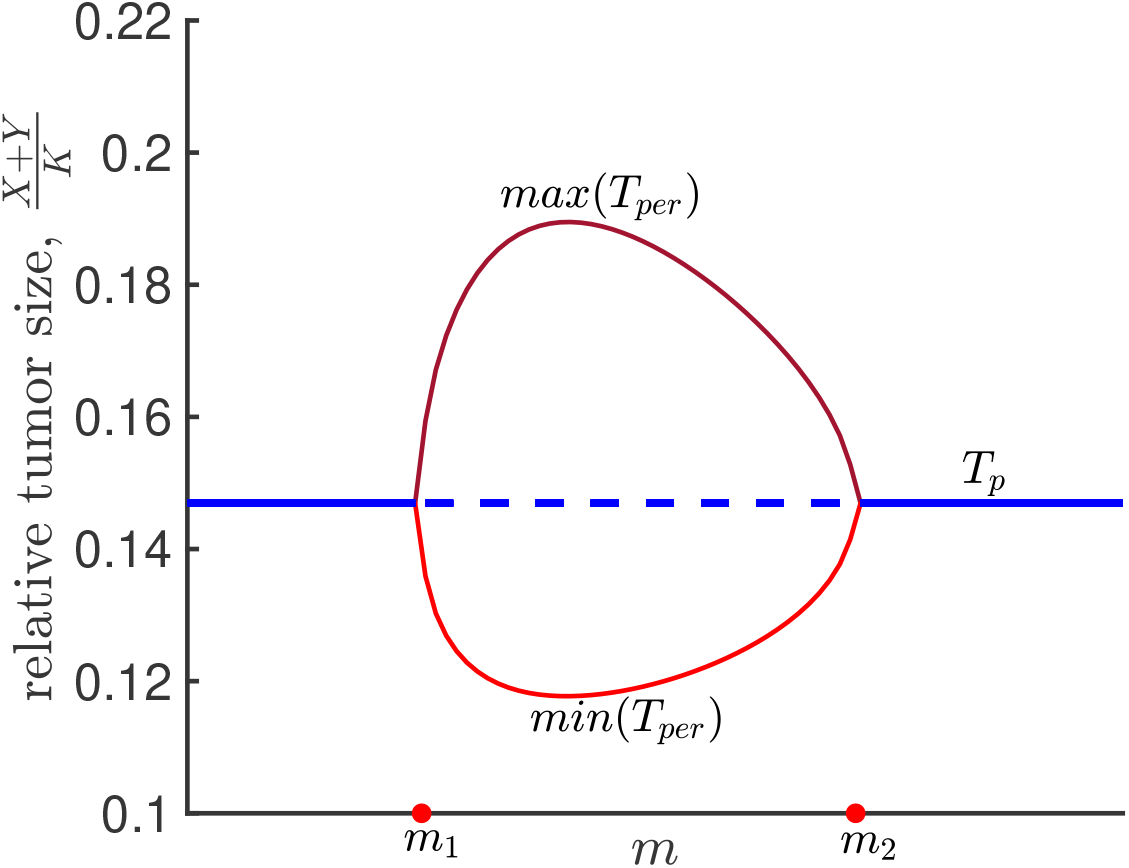
Bifurcation with respect to m: This plot shows how the oscillation in tumor size changes when variable *m* changes. As figure describes when *m* passes through *m*_1_, partially infected equilibrium, 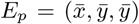 becomes unstable and a stable limit cycle appears. 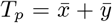, shows the tumor size when *E*_*p*_ is stable, and *max*(*T*_*per*_) and *min*(*T*_*per*_) shows the maximum and minimum of tumor size when the limit cycle exists. When *m* passes through *m*_2_ the stable limit cycle disappears and again *E*_*p*_ becomes stable. This simulation is done for the set of parameters *s*=0.5, *X*_*U*_ =0.9, *Y*_*I*_ =0.08, and *b* ∼ 5.35. This *b* is the mid point of the interval 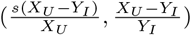, on which *E*_*p*_ is positive. This simulation is conducted for 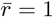, *s* = 0.5, *X*_*U*_ = 0.9, *Y*_*I*_ = 0.08, *m* = 0.1, and *b* 5∼.35. This *b* is the mid point of the interval 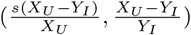, on which *E*_*p*_ exists.

### 5.1 Fast-slow dynamics of tumor size

Another interesting result that is observed numerically is that Model (1) shows fast and slow behavior. As we mentioned above, there is a range of parameters for which there is a unique stable periodic solution to which every trajectory starting from a positive point, except for a single curve (the stable manifold of the steady-state), is attracted. However, not every solution gets attracted with the same speed. Since the speed of remission is a crucial factor in the therapy, it motivates us to investigate in which regions of state spaces fast and slow behavior occur.

Figure 8 which is a modified version of Figure 6 shows fast dynamics. Under the set of parameters that this simulation is conducted equilibrium *E*_*p*_, which is marked with black circle, has three eigenvalues *e*_1_ = −0.212 and *e*2, *e*3 = 0.001 ± 0.11*I*. Therefore, in a neighborhood of *E*_*p*_ solutions oscillate with the period 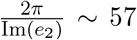. A thousand random points, marked by yellow circles, are chosen to show fast attraction to the global attracting surface. Each solution that starts from a point marked yellow terminates in a purple point after 57 days. In Figure 8 same plot has been provided from two different views to provide a better observation that all the purple points are located on the slow manifold. The slow manifold includes the global attracting surface and extends outside the limit cycle, including the solid green area. Therefore, solutions on the slow manifold tend very slowly to the limit cycle, as we discussed in Figure 1.

**Figure 8:**
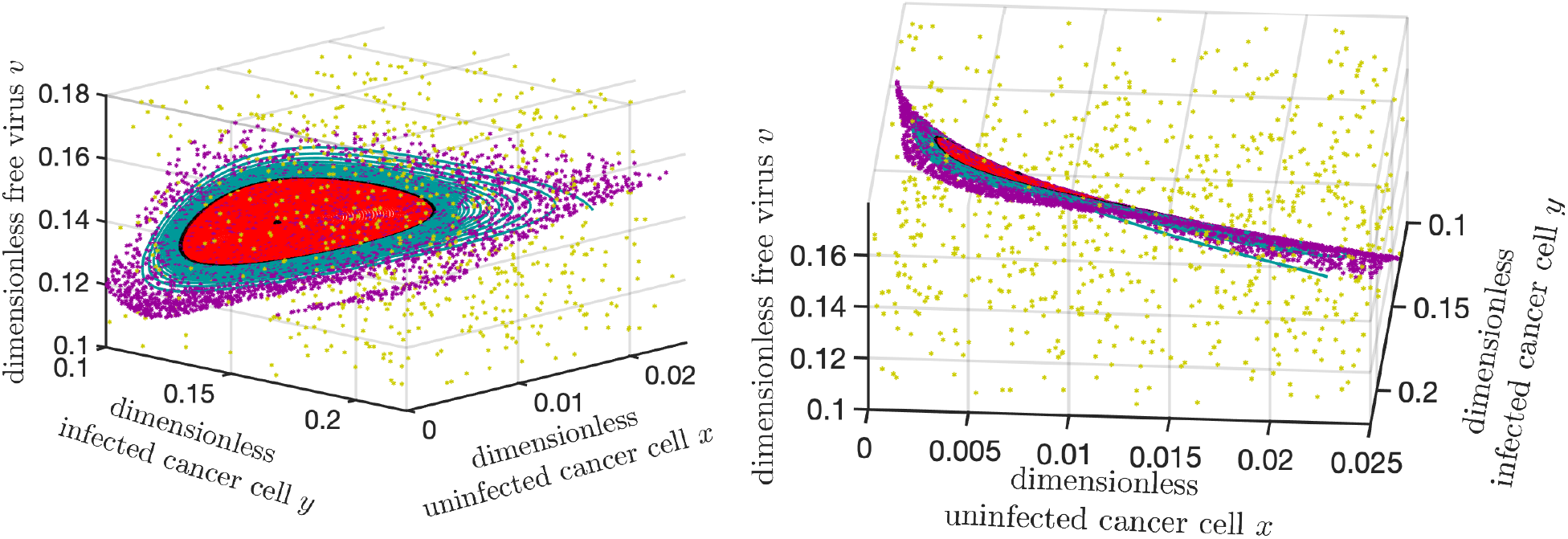
Fast attraction and collapse to the global attracting surface over one period. The two plots are the same simulation shown in different views. The black closed curve bounding the red disk is an attracting limit cycle, and the black circle near the center of the red disk is the equilibrium *E*_*p*_. In this figure, thousand random points, yellow dots, are chosen, and the positions of their trajectories after 57 days are shown with purple dots. All the purple dots appear to lie on a 2-dimensional surface that contains the limit cycle, *E*_*p*_ and the unstable manifold of *E*_*p*_(The global attracting surface is the disk centered at equilibrium *E*_*p*_ and includes the disk boundary, which is the limit cycle). This shows each trajectory tends quickly to the global attracting surface.

An example of slow dynamics is shown in Figure 1, where two solutions are shown approaching a periodic orbit very slowly. Trajectories oscillate with a period of about 57 days. It takes years to get close to the periodic orbit. One (green) approaches with decreasing amplitude, and the other (red) approaches with increasing amplitude. The black curve denotes the equilibrium *E*_*p*_. This is the equilibrium that loses its stability at the Hopf bifurcation point when the stable limit cycle is created. The red curve starts with an initial condition near *E*_*p*_ and slowly spirals out to the stablelimit cycle. The two yellow circles denote the time interval in which the red trajectory’s amplitude grows from 10% of the limit cycle’s amplitude to 90%, a period of 8.2 years. In this Figure 1 we have used the parameter values 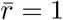, *X*_*U*_ = 0.9, *s* = 0.5, *Y*_*I*_ = 0.08, *m* = 0.1 and *b* 5. ∼ 35. This *b* is the midpoint of the interval 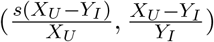, on which *E*_*p*_ exists.

In Figure 9 we use the Poincare map defined on the Poincare surface *y* = *v* to observe where the slow manifold is located. The black curve is the periodic solution and the black triangle is equilibrium *E*_*p*_. The yellow circle on the periodic solution represents the fixed point ℱ 𝒫 of the Poincare map, which in this case it takes 57 days for the Poincare map starting from fixed point to return back to the fixed point. The rest of the curves are obtained by capturing the orbit of a given point under the Poincare map. The red curve and the green curves in this figure show slow dynamics. For example for the red curve it takes at least 80 iterations of the Poincare map for the initial point chosen near to the equilibrium to tend to ℱ 𝒫 and it takes up to 55-57 days between two consecutive points in the orbit, meaning it takes at least 4480 days to approach towards the limit cycle. Starting from any initial condition not located on the green or red curve, 2nd or third point in the orbit under Poincare map is located either on the red or green curve. The Jacobian of the Poincare map at ℱ 𝒫 has two eigenvalues *e*_1_ = 7.3295 · 10^(*−*6)^ and *e*_2_ = 0.8962. Since *e*_2_*/e*_1_ = 122280 means that the fast attraction happens 122280 times faster than the slow dynamics.

**Figure 9:**
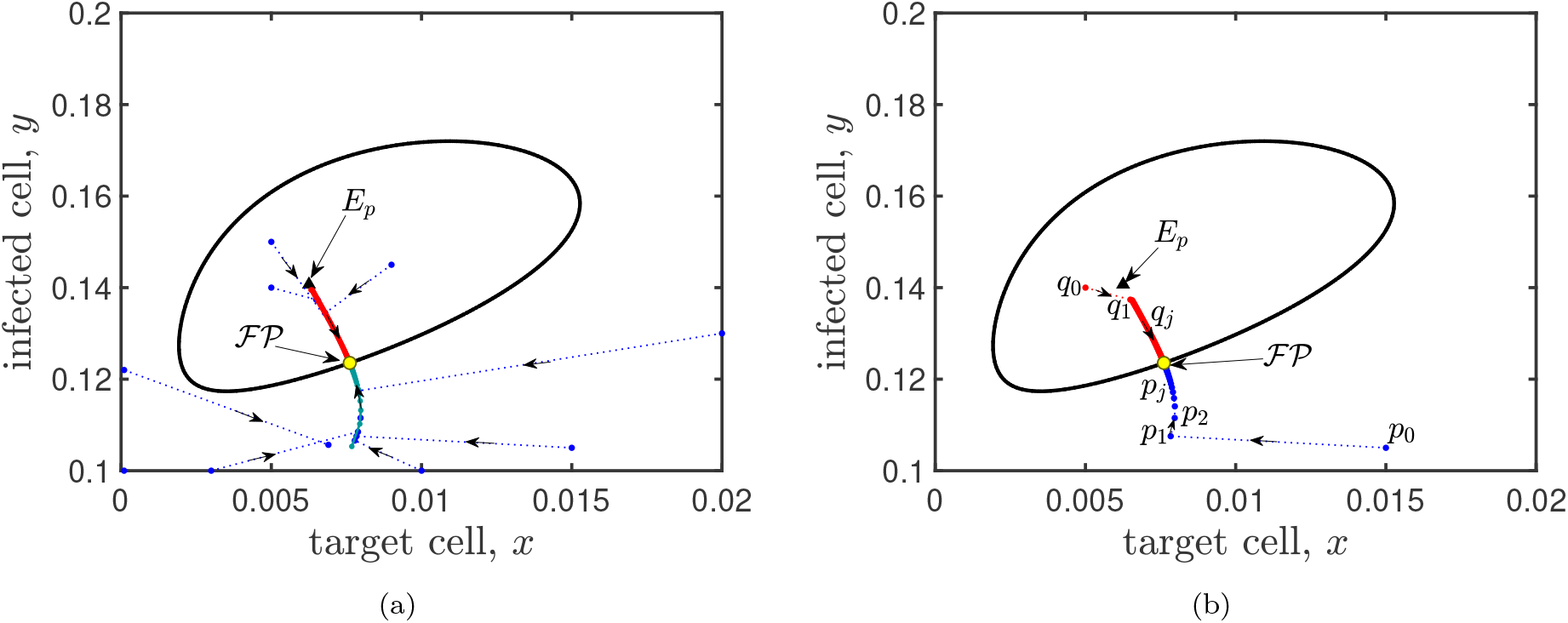
Capturing fast-slow dynamics using Poincare map. **(a)** This plot shows slow attraction to the limit cycle and fast attraction to the slow manifold. Except the black curve which shows the stable limit cycle, every other curve is the positive orbit of a given point under the Poincare map. Every blue curve starts from an initial point which is not located on the red or the green curves. The second or third point in the positive orbit of each given point which is not located on the red or green curve, is located on the red or green curve. This assures fast attraction to the green and red curve. Any point chosen on the green and blue curve takes a lot of iteration to tends to the point marked yellow. The yellow point is on the Poincare section and it returns to itself after one iteration, therefore it is the fixed point of the Poincare map. The set of all the points on the green and red curve except the fixed point of the Poincare map, is called the slow manifold. Plot **(b)** is conducted to show how each orbit of a given point under the Poincare map behaves. To do this, two orbits *q*_*j*_ starting from *q*_0_ and *p*_*j*_ starting from *p*_0_ are shown. Both orbits tend to the fixed point of Poincare map labeled by ℱ 𝒫. The first iteration from *p*_0_ to *p*_1_ takes 55 days, it takes between 56-57 days between every two consecutive points in the orbit of *Pj*. The time difference between every consecutive point in the sequence *q*_*j*_ changes from 60-57 days and eventually remains 57 days. This simulation is conducted for 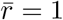, *s* = 0.5, *X*_*U*_ = 0.9, *m* = 0.1 and *b* ∼ 5.35. This *b* is the mid point of the interval 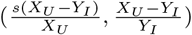, on which *E*_*P*_ exists.

### 5.2 Dependency of virulence threshold on viral time scale and horizontal transmission rate

Recall parameter 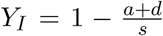. Changes in parameters *a* and *s* lead to changes in *Y*_*I*_. A more aggressive virus causes more increase in death rate and more decrease in the growth rate of cancer cells after infection, i.e., higher *a* and lower *s*. Notice that when *a* increases or *s* decreases, *Y*_*I*_ decreases. Hence, lower *Y*_*I*_ values mean a stronger virus is administrated. Thus, we can use *Y*_*I*_ as a measure of the level of cytotoxicity of the virus. It is important to know how virulent a virus should be engineered to obtain more tumor shrinkage under therapy. Our simulations below suggest that identifying the optimum range for virulence level of virus depends on both the viral time scale and infection horizontal transmission rate.

Figure 10 is a contour plot of maximum tumor size with respect to parameters *b* and *Y*_*I*_ in two scenarios; (a) fast viral time scale, *m* = 1 (b) slow viral time scale, *m* = 0.1. This figure recommends when parameters *b* and *Y*_*I*_ are chosen in the area below the solid curve (this curve is labeled by 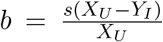, then therapy is not effective regardless of viral time scale. Additionally, when parameters *b* and *Y*_*I*_ are chosen in the area above of the dotted curve (this curve is labeled by 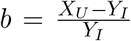, for a fixed *b* value, by decreasing *Y*_*I*_, tumor size decreases, regardless of viral time scale.

**Figure 10:**
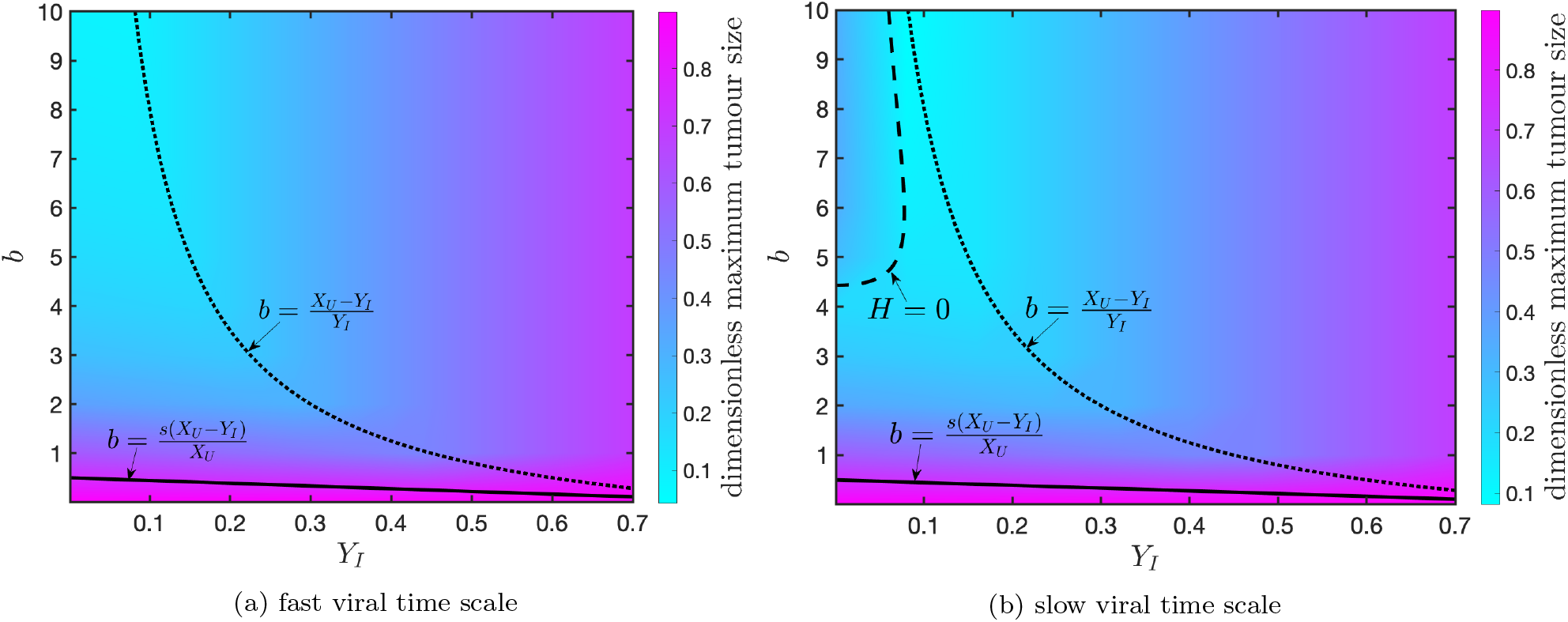
Changes in the maximum tumor size with respect to *b* and *Y*_*I*_. This figure represents how tumor size changes when parameters *b* and *Y*_*I*_ changes. Parameter values are *s* = 0.5, *X*_*U*_ = 0.9, and *m* = 1 in plot (a), and *m* = 0.1 in plot (b). Plot (a) shows when virus dynamics are fast, the maximum tumor shrinkage under therapy is possible for lower *Y*_*I*_ and higher *b* values. Plot (b) shows when virus dynamics are slow, lower *Y*_*I*_ values will not guarantee the best outcome. In this case, the smallest tumor size is achieved when parameters *b* and *Y*_*I*_ are chosen such that *H* = 0.

Figure 10 suggests the viral time scale affects the outcome of therapy when parameters *Y*_*I*_ and *b* are chosen in the area between the solid and dotted curves.

As Figure 10a illustrates, when the viral time scale is fast, the best range for parameter *b* and *Y*_*I*_ is the area between the solid curve and the dotted curve.

When the viral time scale is slow, as Figure 10b represents, the most tumor shrinkage is obtained when the parameters *Y*_*I*_ and *b* are chosen along the dashed curve. The dashed curve labeled by *H* = 0 shows an implicit plot of roots of the Routh function (Eq. (13)). Assume *b* = 5.35 is fixed, in this case *H*(*Y*_*I*_ = 0.08) = 0. When *b* = 5.35 is fixed and *Y*_*I*_ decreases and passes through the value 0.08, the maximum tumor size increases. Therefore, Figure 10b suggests when viral timescale is slow, a high level of virulence has drawbacks on the outcome of viral therapy. Therefore, when the viral time scale is slow, the best range for parameters *Y*_*I*_ and *b* is the area between the three curves. In Figure 11, five different values of *Y*_*I*_ are chosen to find out the effective level of virus cytotoxicity for a fixed *b* value. The value of *Y*_*I*_ for which the red curve is simulated is 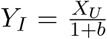. The value of *Y*_*I*_ for which black curve is plotted, when *m* = 0.1, is the value of *Y*_*I*_ for which *H* = 0.

**Figure 11:**
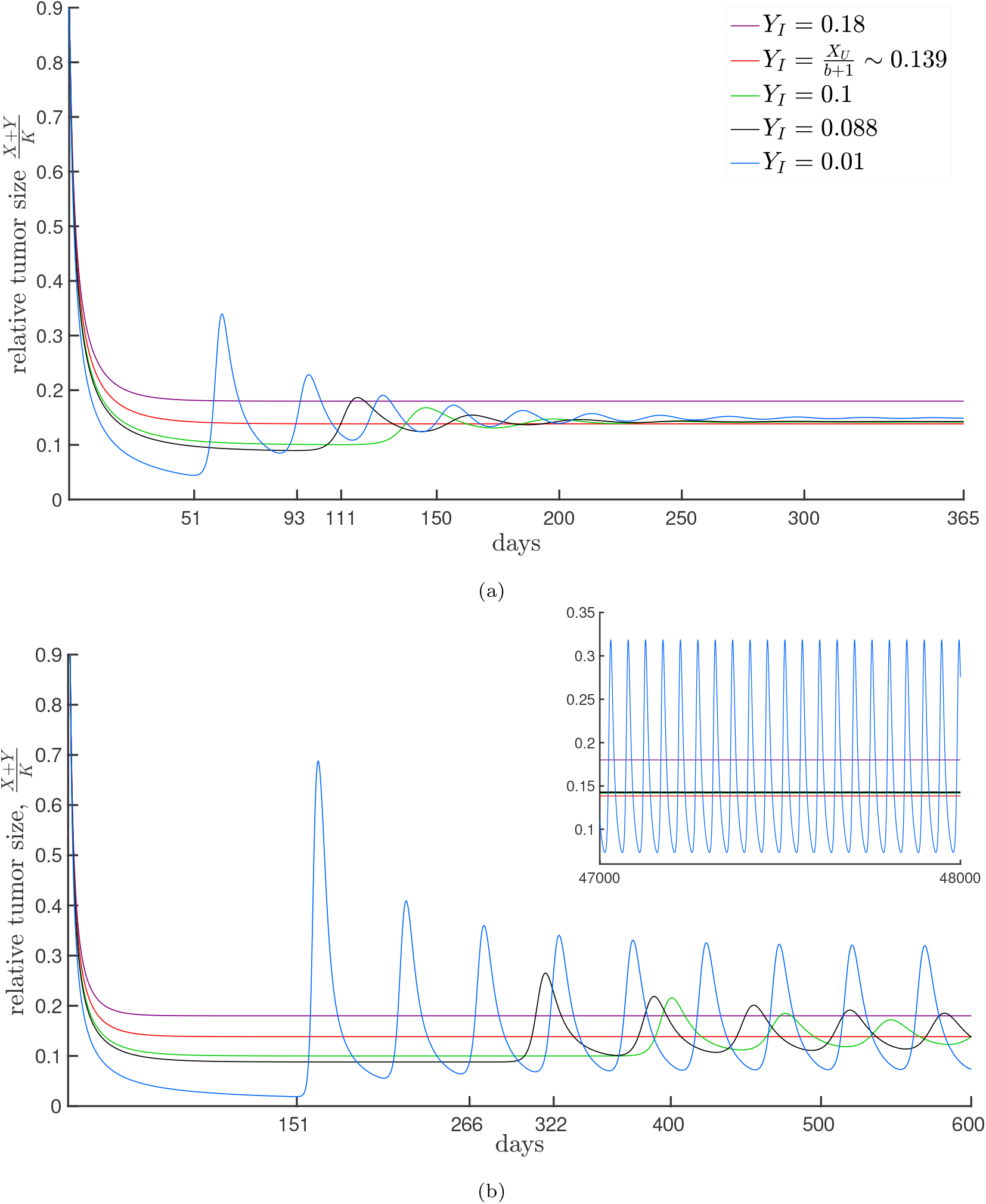
Effect of cytotoxicity *Y*_*I*_ on the tumor size. Plot (a) *m* = 1 and Plot (b) *m* = 0.1. As Plot (a) shows for higher values of *m*, when 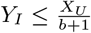, eventually the tumor size is smaller. Therefore, the best range for parameter *Y*_*I*_, 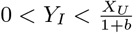. For *m* = 0.1, as plot (b) suggest when values of *Y*_*I*_ decrease, dynamics becomes oscillatory and the amplitude of oscillation increase by decreasing *Y*_*I*_. For *Y*_*I*_ = 0.01 stable oscillation occurs with a very high amplitude. As the small panel at the right top corner of plot (b) shows, both green and black curve or decaying oscillation but it takes very long time for the amplitude of the black curve to tends to zero. When *m* = 0.1, the best range for 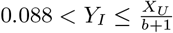. This simulation is conducted under the set of parameters 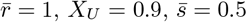, and *b* = 5.5.

In the case fast viral time scale, in other words when *m* is high enough that it is not in the interval (*m*_1_, *m*_2_), no periodic orbit exists. In this case as Figure 10a and Figure 11a suggest for 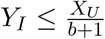, changes in *Y*_*I*_ does not affect the tumor size much. The blue, black and green curve in Figure 11a are all corresponding to the cases where 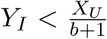. There is not much difference in their eventual tumor size with the case when 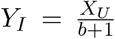. However, the tumor size is eventually smaller when 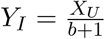. Therefore, for higher values of *m*, and fixed *b* value, the best range for *Y*_*I*_ is 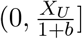 and the most tumor shrinkage is achievable at 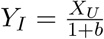.

Figure 11b shows the tumor size overtime when the viral time scale is slow and parameter *b* is fixed. As the figure illustrates, when 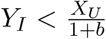, the system shows oscillatory behavior: both green and black curves are decaying oscillations. The value of *Y*_*I*_ for which the black curve is simulated is when *H* = 0, therefore a Hopf-bifurcation occurs at this point, *Y*_*I*_ = 0.088. Therefore, the amplitude of the oscillations takes a very long time to tend to zero. Hence, in practice, if solutions are only observed for a short time, the black curve seems like a stable periodic solution. When *Y*_*I*_ = 0.01 *<* 0.088 (the blue curve), the tumor size is periodic with a very high amplitude. As the small box in the right corner of Figure 11b suggests, there is not much difference in the eventual tumor size for the black, green and red curves. If the solution is observed for a very long period of time, eventually, the red curve would be lower, but in practice, it is not possible. Therefore, when the viral time scale is slow, for a fixed *b* value, the best range for *Y*_*I*_ is between the value of *Y*_*I*_ at which *H* equals zero and 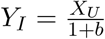. In this case, the most tumor shrinkage is observed at the value of *Y*_*I*_ for which *H* = 0.

## 6. Discussion

This paper offers a very simple mathematical model, Model (1), to analyze the interactions between cancer cells and free viruses during an oncolytic viral therapy. Even though Model (1) is simple, it captures some important features about oncolytic viral therapy: features that help to set expectations about the outcome of the treatment and how to engineer the vaccine in order to get better results. For example, Lemma 3 shows the tumor eradication equilibrium, *E*_0_, is always unstable. This means complete tumor eradication is not possible. This suggests an oncolytic virus that only has an oncolysis effect cannot be used as a cure for cancer. However, Proposition 1 implies that the tumor size under therapy over time is always smaller compared to the case with no therapy. Therefore, oncolytic viral therapy can be used to control the speed of growth of a tumor. Proposition 2 and Lemma 3 together show that oncolytic viral therapy can be used to reduce the tumor size considerably. The proposition suggests oncolytic viral therapy may be used as an intervention method to control the size of the tumor before a conventional therapy can be followed up.

One of our goals in this study was to optimize the therapy by recommending the parameter regions in which the therapy has the best outcome. To do this, we focused our analysis on some parameter combinations: a measure of virulence level, *Y*_*I*_, a rescaled horizontal transmission rate, *b*, and the viral clearance rate relative to the mutation rate of uninfected cancer cells, *m*.

To ease the communication with the reader throughout the paper, we denoted the stable, steady state tumor size when therapy fails, or if no treatment occurs, by *X*_*U*_ and the minimal steady state tumor size under the therapy by *Y*_*I*_. In Proposition 2, we showed that the interval [*Y*_*I*_, *X*_*U*_] is a global attracting set for the tumor size. Specifically, for a specific virus, i.e., for a fixed set of parameters, the tumor size by therapy is eventually bigger or equal to *Y*_*I*_. Therefore, *Y*_*I*_ is the minimum tumor size under the therapy.

Lemma 3 and Theorem 2 specify two possible outcomes for the treatment: 1-failure, 2-partial success or tumor control. Partial success means a considerable reduction in tumor size. Partial success includes two scenarios: 2(a) tumor partially gets infected 2(b) 100% infection prevalence occurs.

### 1-Failure

Treatment fails when the horizontal rate of infection transmission is smaller than the control threshold. In this case, therapy has no impact, and the tumor grows to the maximum size *X*_*U*_, the same as when no therapy would take place. Long-term tumor reduction will be possible when the horizontal rate of infection transmission is above of control threshold.

### 2.-Partial success

#### 2(a). partial infection

This scenario occurs when the infection transmission rate is between the control threshold and optimum threshold. In this case, there are two possible scenarios; if the viral time scale is slow, the positive equilibrium is unstable, and every solution starting from a positive point that is not on the stable manifold of the positive equilibrium tends to the stable limit cycle. When the viral time scale is fast, the positive equilibrium is globally stable, and no limit cycle exists. See Figures 6 and 7. In this case, as Propositions 1 and 2, and Lemma 2 suggest since horizontal infection transmission is above the control threshold, therapy reduces the tumor size, in comparison with the case that no therapy occurs, but it is not the minimum possible tumor size under the therapy.

#### 2(b). 100% infection prevalence

The maximum tumor shrinkage by therapy is obtained in this scenario which happens if infection horizontal transmission is equal to the optimum threshold.

In Model (1), we assumed (assumptions (A2) and (A4)) cancer cells are mitotic, and the growth rate of infected cancer cells is lower than their apoptosis rate. If we relax this assumption, meaning either the infected cells do not self-renew or the self-renewal rate of infected cancer cells be lower than their apoptosis rate, then 100% infection prevalence will not be possible. Therefore, only two scenarios would be possible: failure or partial infection. Furthermore, the tumor shrinkage by therapy will be less than our model, which considers two pathways to infection transmission.

In this study, interactions with immune cells and spatial heterogeneity (neither morphological nor phenotypical) are not discussed. Model (1) can be extended to consider the effect of virus specific immune response that seems to be one of the main barriers in the success of oncolytic viral therapy. In our work in progress, by extending the model to also consider interactions of virus specific immune responses with cancer cells and the free virus, in most cases, no longer oscillatory behavior is observable. In the presence of virus specific immune responses, the reduction in tumor size will be lower in comparison with the case no interaction with immune cells is taken into account. However, oncolytic therapy seems yet promising as a considerable reduction in tumor size is still observable.

One of the assumptions under which we started the study was assumption (A5). One might question how virulent the virus should be? As we discussed before, parameter *Y*_*I*_, the minimum tumor size, can also be used as a measure of the level of cytotoxicity of the virus. Lower values of *Y*_*I*_ mean the virus is more virulent. As Figures 10 and 11 illustrate electing the best range for virulence level also depends on the parameter *m*. Parameter *m* is the relative clearance rate of the free virus and gives the timescale of the virus dynamics. When parameter *m* is large, meaning 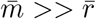, as earlier studies by Komarova and Wodarz [18] suggested the system is in quasi-steady state. Meaning, *y* −*v* rapidly approaches zero, therefore Model 1 can be reduced to the “slow” *x*-*y* subsystem. The maximum level for cytotoxicity recommended by Komarova and Wodarz [18] in terms of parameters of Model (2) is equivalent to 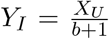 (notice that maximum level of cytotoxicity means lowest value recommended for *Y*_*I*_). However, our results show that if the parameter *m* is high enough that does not belong to the interval (*m*_1_, *m*_2_) (see Theorem 2 for *m*_1_, and *m*_2_), then 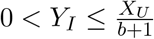. Our simulations in Figures 10 and 11 recommend the highest value for *Y*_*I*_ is *X*_*U*_ */*(*b* + 1). Although, there is not a huge difference in the amount of tumor shrinkage corresponding to *Y*_*I*_ values within the interval (0, *X*_*U*_ */*(*b* + 1)). If *m* ∈ (*m*_1_, *m*_2_), then *Y*_*I*_ should be chosen between the values of *Y*_*I*_ for which the Routh function *H* = 0 (see Eq. (13)), and 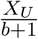. This suggest when *m* is lower, in other terms, virus dynamics is slow, a very high level of virulence is not recommended.

As we mentioned above, oncolytic viral therapy can be used as a method for controlling tumor size. What we suggested above about parameter regions is mainly considered when the viral therapy is going to be used alone. Therefore, we recommend the parameters are chosen in a way that sustained oscillations do not occur. Specifically, parameters should be chosen in the way *H* ≥ 0. In contrast, if oncolytic viral therapy is going to be followed by another therapy, one may reconsider the lower values of *Y*_*I*_ for which sustained oscillations occur. As we observed in Figure 11b, when the virus is highly virulent and the proper initial dosage of the virus is applied, early tumor shrinkage occurs to less than 11% of the initial tumor size, and the tumor remains in this size for almost a year before starting to relapse. For example, when *Y*_*I*_ = 0.088 (the black curve), tumor size reaches to 9.8% of initial tumor size at *t* = 266 or the most aggressive case when *Y*_*I*_ = 0.01 tumor size reaches to 2% of initial tumor size at *t* = 150. This raises multiple questions, such as, what happens if the treatment switches to another therapy before tumor relapse? Which treatment would be best as the followed-up treatment? We suggest that a virulent virus can be used, but before the tumor relapses, the second therapy must be followed up.

## Appendix A. Derivation of dimensionless Model (2)

Here we derive the dimensionless Model (2). First we recall Model (1).

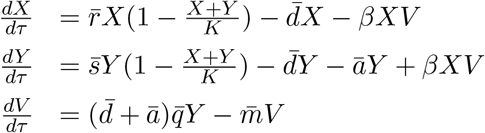

Now let 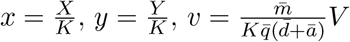 and 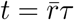, then

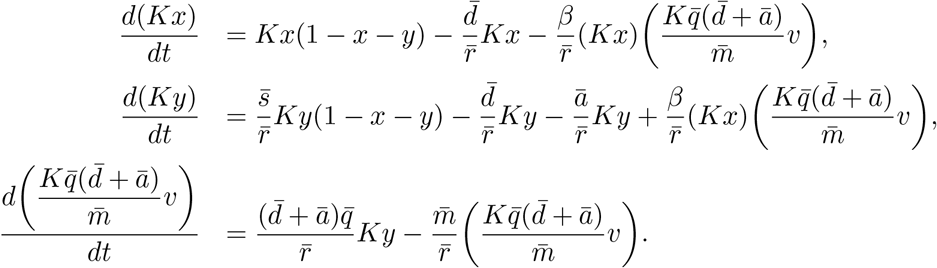

After some algebraic simplifications the above system of equations turns to

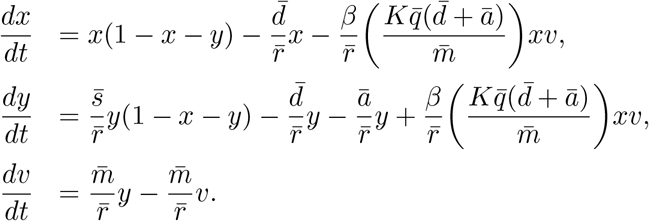

Now we will call dimensionless parameters as follows:

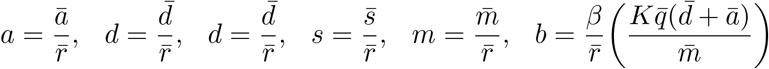

Using the new dimensionless parameters the above system of equation gets the following form.

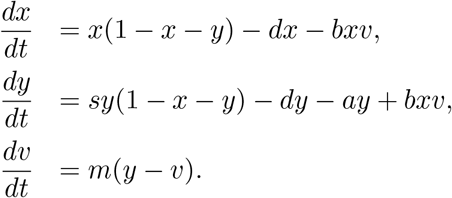

Now let *X*_*U*_ = 1 −*d* and 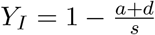, then Model (1) gets the following dimensionless form.

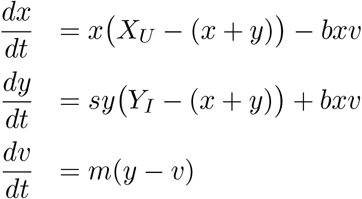

## Appendix B. Detailed proof of lemma 1

In this section we will show what happens in each boundary of the set W.

On the *x* axis Model (2) simplifies to: 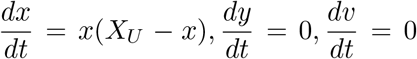. Hence, each solution starting from a point on the *x* axis with *x* ≥ 0 remains on the *x* axis. Except *E*_0_ = (0, 0, 0) which is an equilibrium of the system, each solution (*x*(*t*), 0, 0) with *x*(0) *>* 0 tends to *E*_*U*_ = (*X*_*U*_, 0, 0) as *t* increases.

On the *y* axis Model (2) simplifies to: 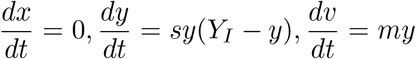. Since 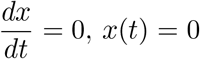, *x*(*t*) = 0 for all *t* ≥ 0.

The solution to ode 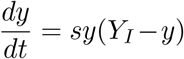, with initial point *y*(0), is 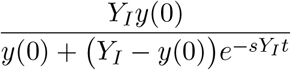. Therefore, if *y*(0) *>* 0, then *y*(*t*) *>* 0 for all *t* ≥0.

Since 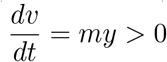 when *y >* 0, *v* is increasing. Therefore, *v*(*t*) ≥ 0 for all ≥ 0.

Hence, each solution starting from a point (0, *y*_0_, 0) with *y*_0_ *>* 0 remains on the *y* − *v* plane.

On the *v* axis Model (2) simplifies to: 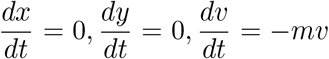. Therefore, a solution starting on the *v* axis, remains on the *v* axis. Additionally, since 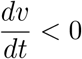 when *v >* 0 each solution starting from a point on the *v* axis with *v*_0_ *>* 0, tends to *E*_0_.

On the *x*-*v* plane *y* = 0. Hence, Model (2) simplifies to: 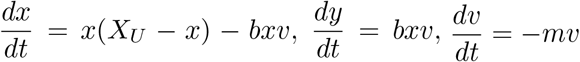.

Therefore, *v*(*t*) = *v*(0)*e*^*−mt*^. *v* is non-negative and decreasing. Since 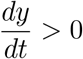 when *x, v >* 0, starting from an initial point with *x*(0), *v*(0) *>* 0, *y* increases. Therefore, *y* remains positive. Thus, any solution starting from a point in the *x*-*v*, except the points on the *x* and *v* axes, tends inward *W*, i.e. in the direction of increasing *y*.

On the *x*-*y* plane *v* = 0. Thus, Model (2) simplifies to; 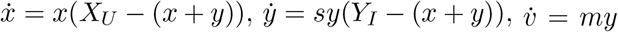. When *y*(0) *>* 0, 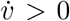. Hence, *v* is increasing. Each solution that starts from a point (*x*(0), *y*(0), 0) ∈ 𝒲 where *y*(0) *>* 0, points inward the 𝒲 as *t* increases.

On the *y*-*v* plane, *x* = 0. Thus, Model (2) simplifies to: 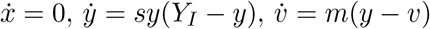.

Since 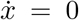, therefore *x*(*t*) = 0 for all *t* ≥ 0. Hence, each solution starting on this plane, is tangent to the plane. Solution to the foregoing system of differential equations with an initial point (0, *y*(0), *v*(0)) is

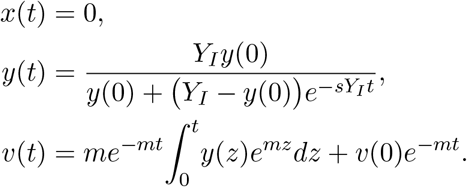

Therefore, each solution that starts from a point in the *y*-*v* plane with *y*_0_ ≠ 0 goes eventually to *E*_*I*_. Figure B.12 shows the vector field on the y-v plane.

**Figure B.12:**
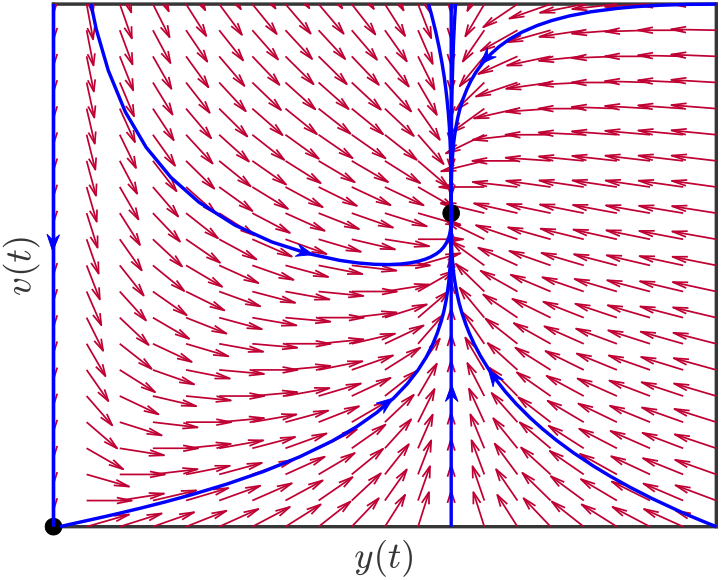
Any solution that starts in the *y*-*v* plane will remain the y-v plane. All the solutions starting in *y*-*v* plane with *y*(0) *>* 0, eventually tends to *E*_*I*_.

The intersection of 𝒲 with the plane *v* = *X*_*U*_ is above of *y* = *v* plane, this boundary only meets the plane *y* = *v* at point (0, *X*_*U*_, *X*_*U*_)). Hence, 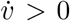 *<* 0 every where on this boundary except point (0, *X*_*U*_, *X*_*U*_). Therefore, *v* is decreasing. Therefore, any solution starting on this plane points inward of 𝒲. Since (0, *X*_*U*_, *X*_*U*_) is located in *y*-*v* plane, the solution starting from (0, *X*_*U*_, *X*_*U*_) tends to *E*_*I*_.

When 𝒲 meets the plane 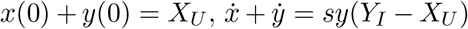. According to assumption (H2), *Y*_*I*_ *< X*_*U*_. Thus, *x* + *y* is decreasing when *y* ≠ 0, so each solution that starts from a point on the intersection of 𝒲 with plane *x* + *y* = *X*_*U*_, points inward 𝒲.

## Appendix C. Hopf Bifurcation

Consider a three dimensional system of ordinary differential equations is given, meaning

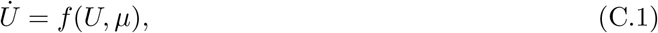

where *U* ∈ ℝ^3^ and *μ* ∈ ℝ.

**Hopf Bifurcation Theorem**. A Hopf-bifurcation occurs at (*U* ^*\**^, *μ*^*\**^) for the system (C.1) if there exists a curve of stationary points (*U*_*eq*_(*μ*), *μ*) ∈ ℝ^3+1^ such that (*U*_*eq*_(*μ*^*\**^), *μ*^*\**^) = (*U* ^*\**^, *μ*^*\**^) and *Df* (*U*_*eq*_(*μ*) *μ*), has a pair of complex eigenvalues *λ*(*μ*) = *R*(*μ*) ± *iI*(*μ*), such that

HB(1) *R*(*μ*^*\**^) = 0 and *I*(*μ*^*\**^) ≠ 0,

HB(2) 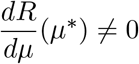, and the real eigenvalue of *Df* (*U*_*eq*_(*μ*) *μ*), is nonzero for all values of *μ*.

### Appendix C.1. Comparison Theorem

#### Theorem 3

(First Comparison Theorem [31]). Let *x*(*t*) and *y*(*t*) be continuously differentiable functions from [0, +∞) to ℝ, satisfying

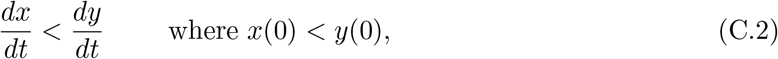

then, *x*(*t*) *< y*(*t*) for all *t >* 0.

## Acknowledgments

We thank James Yorke and Kamran Kaveh for their helpful comments. James Watmough was partially supported by NSERC (RGPIN-2017-05760) and CIHR, the Mathematical Modelling of the COVID-19 Task Force. Lin Wang was partially supported by NSERC (RGPIN-2020-04143). James Watmough and Lin Wang were also partially supported by New Brunswick Health Research Fund.

## Notes

### Competing Interest Statement

The authors have declared no competing interest.

